# Human precentral gyrus neurons link speech sequences from listening to speaking

**DOI:** 10.1101/2024.12.27.630563

**Authors:** Duo Xu, Jason E. Chung, Alexander B. Silva, Sean L. Metzger, Quinn R. Greicius, Yizhen Zhang, Matthew K. Leonard, Manish K. Aghi, Shawn L. Hervey-Jumper, Edward F. Chang

## Abstract

Speech perception and production are interconnected processes, but the underlying neural mechanisms remain unclear. We investigated this relationship by recording large-scale single-neuron activity in the human brain during a delayed sentence repetition task. Contrary to the traditional view that the precentral gyrus is solely responsible for motor execution, we found that neurons there encoded activity across all task phases of listening, delay, initiation, and speaking. Notably, we discovered “mirror” neurons that activated transiently after hearing and before producing specific speech sounds, and “bridge” neurons that maintained activity between the same speech elements during listening and speaking. Population analysis revealed distinct latent components for each task phase and persistent dynamics for specific sentences. Overall, this study provides novel insights into the neuronal basis of speech processing, emphasizing the intricate interplay between perception, production, and verbal working memory.

## Introduction

Fluent speech is one of humanity’s most remarkable abilities, enabling us to share thoughts, express emotions, and build complex social connections. At its core, speech involves a delicate interplay between hearing and vocalizing: the brain must not only perceive and interpret auditory input but also orchestrate precise movements of the vocal tract to produce sound. This ability to transform what we hear into what we say is essential for vocal learning—a process that allows us to acquire and refine language. Through speech repetition, the brain establishes crucial links between sensory and motor representations, forging the pathways that make fluent communication possible ^1^. However, this process demands more than just hearing and speaking; it requires a robust short-term memory system to hold auditory information long enough for it to be reproduced. Despite decades of research into the brain areas involved in speech perception, production, and memory, the exact neural computations that integrate these processes remain a mystery, leaving open key questions about how the brain transforms sound into speech.

While extensive research has mapped the brain regions involved in speech perception, production, and memory, the precise neuronal computations that integrate these processes remain poorly understood. Bridging these domains at the neuronal level is essential for understanding how the brain orchestrates complex speech behaviors.

One region that has garnered significant attention in this context is the human precentral gyrus (PrCG). Traditionally recognized for its role in motor control, recent evidence suggests that the PrCG also plays a pivotal role in speech processing. This area houses multiple functions including vocal-tract articulation ^2–5^, auditory perception ^6,7^, and even reading ^8^. Notably, the middle PrCG has been implicated in phonological motor planning ^9^ and verbal working memory ^10^. However, despite these insights, most studies rely on macroscopic methods such as electroencephalography (EEG) or mesoscopic approaches like functional magnetic resonance imaging (fMRI) and electrocorticography (ECoG). These methods, though valuable, lack the resolution to disentangle the diverse neuronal populations involved in speech functions at the cellular level.

Understanding how speech-related processes are organized within the PrCG, particularly whether functions such as listening, speaking, and verbal memory are integrated or operate independently at the cellular level, requires high-resolution recordings. Specifically, examining how sequences of speech elements are maintained across different task phases can shed light on the underlying neural mechanisms.

In this study, we address these gaps by leveraging high-density Neuropixels probes ^11–13^ to record the activity of 1,794 single neurons across key speech-related regions. These include the PrCG, the pars opercularis of the inferior frontal gyrus (IFG-op, part of Broca’s area), and the auditory speech cortex within the superior temporal gyrus (STG). By employing a delayed sentence repetition task, we aimed to capture neural dynamics across distinct phases: listening, memory maintenance (delay), initiation, and speech production.

Our findings reveal a nuanced functional architecture within the PrCG. We identified neurons that were active in each of the task phases, including “mirror” neurons, which responded during both listening and speaking, and “bridge” neurons, which maintained activity across the delay period, linking the perception and production of the same speech elements. These findings underscore the PrCG’s integrative role in coordinating speech perception and production through shared representations.

Moreover, population-level analyses revealed distinct latent components corresponding to each task phase and persistent sentence-specific dynamics within the middle PrCG. Notably, PrCG neurons exhibited a preference for encoding higher-order phonemic sequences over basic acoustic or articulatory features, emphasizing its role in processing complex linguistic information.

In sum, our results challenge traditional models of speech processing and offer new insights into the neural mechanisms that support the perception, production, and memory of speech. By highlighting the integrative functions of the PrCG, this study provides a framework for understanding how the brain coordinates sensorimotor processes to facilitate seamless speech communication.

### Precentral gyrus neuronal activity tiles the phases of speech processing

Neuropixels recordings were carried out across four key speech centers of the human brain (Figure 1A): the PrCG (middle and ventral) ^9,14^, pars opercularis (posterior Broca’s area), and STG. Recording sites were selected within areas that were later resected as part of the surgical treatment for brain tumor or epilepsy during awake mapping surgeries (Methods). A total of 1794 single-units and 276 multi-units were isolated from 20 cortical insertions (Figures S1A-C). Based on waveform characteristics, we identified 681 putative pyramidal units, 405 putative fast-spiking units, and 984 units with positive spike waveforms (Figures S1D-G) across the cortical depth (Figure S1H).

**Figure 1.**
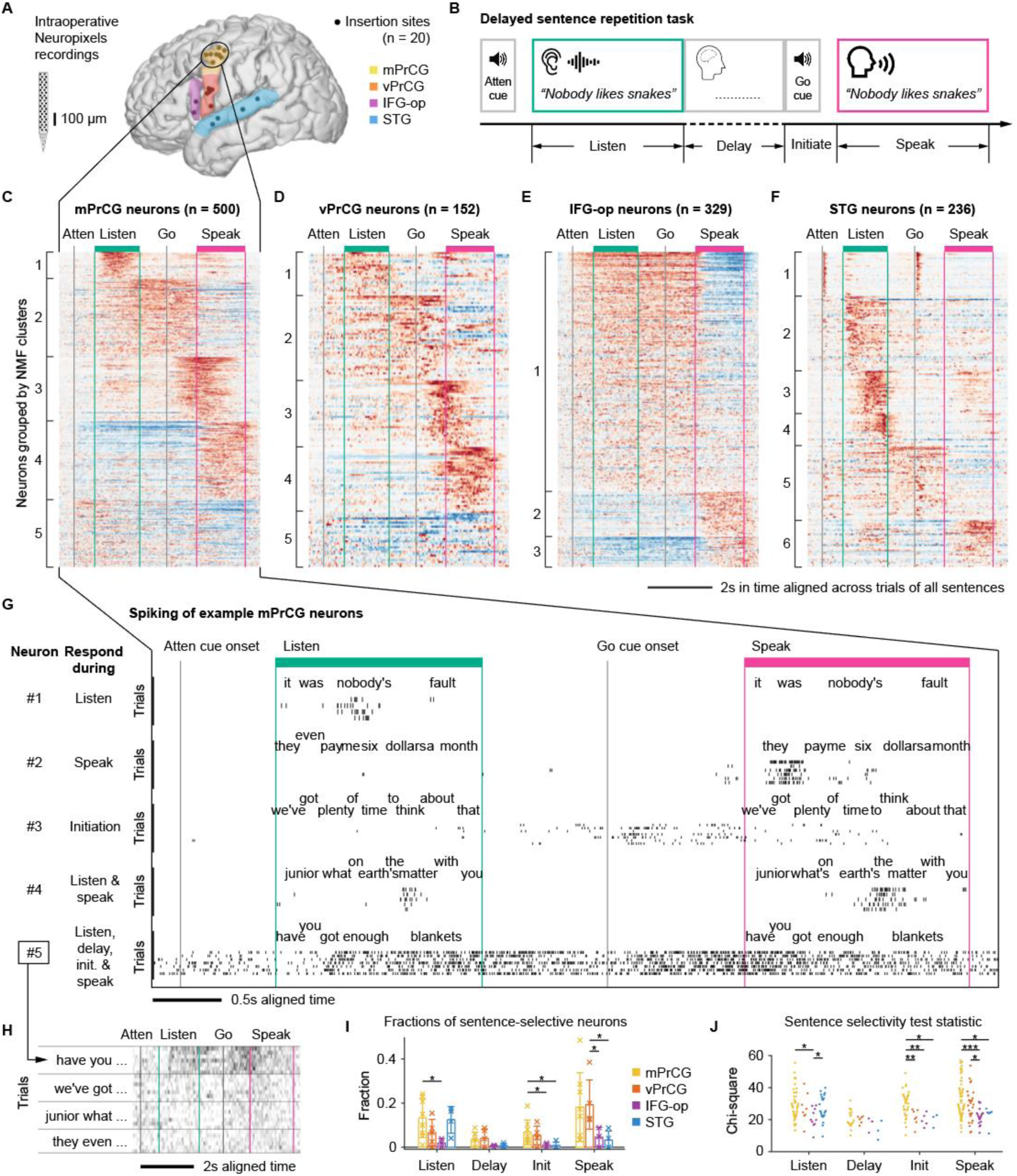
Neuronal activities in precentral cortical columns tile the phases of speech processing. (A) Recording sites and the four regions of interest. (B) Schematic of the delayed sentence repetition task. (C) Average evoked activity of all task responsive mPrCG neurons grouped by NMF cluster membership and sorted by decreasing loadings. (D-F) Similar to (C) but for neurons in vPrCG, IFG-op, and pSTG, respectively. (G) Time aligned and reconstructed single-trial spike rasters of five example neurons from mPrCG, each responds to a specific set of task phases. (H) Single-trial spike rates (rows) of neuron #5 in (G), grouped by four different sentences. (I) Fractions of task-responsive neurons that are sentence-selective in each region (colors) and task phase (groups). Crosses mark fractions of individual recordings. Bar plots show the mean ± SD across recordings. Two-sided Wilcoxon rank sum test. * p < 0.05. (J) The strength of sentence selectivity for each neuron (dot) quantified using the selectivity test statistic (chi-square). Same color and grouping conventions as in (I). Two-sided Wilcoxon rank sum test. * p < 0.05, ** p < 0.01, *** p < 0.001.

Fifteen participants (Table 1) performed a delayed sentence repetition task (Figure 1B; Methods), designed to investigate neuronal activity during four specific phases – listening, delay, initiation, and speaking, as well as how speech content is linked between listening and speaking. In each recording session, we aimed to collect a total of 72 trials consisting of 14 distinct sentences (where 4 sentences occurred 8 times each and the remaining 10 sentences occurred 4 times each), with the goal of sampling the phonetic space of natural speech (Figure S1I) with multiple occurrences of each sentence. Average recording block was 8.8 ± 2.1 min (mean ± SD) and the time was limited given the intraoperative conditions. Task responsive neurons with significant increases in firing rate at one or more task phases relative to baseline (Methods) accounted for 54.1% of all the isolated units in mPrCG, 52.5% in vPrCG, 41.6% in IFG-op, and 56.7% in STG.

To understand when neurons responded during the delayed sentence repetition task, we plotted the task-evoked neuronal activity averaged across trials of all sentences and applied unsupervised (NMF) clustering separately in each region to uncover temporal motifs of neuronal response patterns (Figures 1C-F; Movie 1; Methods). In mPrCG (Figure 1C) and vPrCG (Figure 1D), though a large group of neurons (cluster 3 and 4) show the strongest activity during speaking, other groups responded during listening (cluster 1) and the delay (cluster 2). Similar heterogeneity of single neuron response patterns was observed within individual cortical columns (Figures S2E,F).

Within the temporal motifs highlighted by the clustering, neurons in mPrCG and vPrCG can show significant activity during more than one task phase (Figures S2A-B). To examine the diversity of PrCG neuronal activity, we identified example neurons with distinct task phase specific response profiles. The first example PrCG neuron (Figure 1G, neuron #1) was active only during listening and responded in the middle of “*nobody’s*”. Neuron #2 was active only during speaking, particularly before the utterances of “*pay*”. We also found neurons, such as #3, which showed the strongest activity during speech initiation. More surprisingly, we observed neurons with highly specific firing patterns in both listening and speaking, such as neuron #4. In addition, there were neurons that were persistently active across task phases from listening through speaking, such as neuron #5. The persistent activation of neuron #5 was only present in the sentence “*have you …*” but not the other three (Figure 1H). These last two examples indicate a role for individual PrCG neurons in the multiple sensorimotor processes required for the delayed sentence repetition task. Across all task-responsive neurons, 14.7 ± 3.8% in mPrCG and 13.7 ± 8.4% in vPrCG are significantly sentence-selective (Kruskal-Wallis test; Methods). These PrCG neurons demonstrate distinct task phase-speech content-specificity.

In contrast to PrCG, IFG-op neurons demonstrated minimal selectivity during the task phases and across stimulus. Most IFG-op neurons showed uniform average activity during the entire period from listening to delay to initiation (Figure 1E, S2G; cluster 1) with the remaining groups responding during speaking (cluster 2 and 3). The proportions of sentence-selective IFG-op neurons were small and significantly less than PrCG during listening, initiation, and speaking, as quantified in Figure 1I (comparing purple with yellow and/or red). IFG-op neurons were generally not selective to sentences, especially compared to PrCG, as quantified in Figure 1J (comparing purple with yellow and/or red). For example, we plotted single-trial responses of the three neurons with quantitatively strongest selectivity in each region (Figure S2I-L). The three IFG-op neurons did not show differential sentence responses as clearly as those in other regions.

The majority of STG neurons (Figure 1F) showed the strongest activity when listening to speech (cluster 2-4), consistent with prior work ^15^, or to auditory cues (cluster 1). We also found a smaller group of neurons most active during speaking (cluster 6). The proportion and strength of sentence-selective STG neurons was comparable to PrCG during listening (Figure 1I,J; n.s.). However, very few STG neurons were sentence-selective during speaking.

Together, we demonstrated the specific neuronal activation patterns across the cortical speech network. PrCG contained a functionally integrated local circuit that carried speech information in diverse task phases of listening, delay, initiation, and speaking. Subgroups of neurons in this local circuit further unpacked this integrated function with firing to distinct combinations of task phases. In contrast, IFG-op neurons minimally distinguished task phases or the speech content, and therefore did not support a role of IFG-op in speech planning or articulation. STG neurons signaled speech information primarily during listening and were suppressed during speaking, presumably due to feedback of self-voice.

### “Mirror” and “bridge” neurons link speech elements from perception to production

A central question in speech repetition is the mechanism that links the heard speech with later spoken utterances. Our findings of single neurons that responded to multiple task phases across listening and speaking (e.g., Figure 1G, #4,5) point to a potential solution. We wondered whether these neurons were activated by related speech content and linked the representations from one phase to another.

When we examined the response timing to individual sentences, we observed that firing was highest during similar parts of the sentence in both task phases (Figures 2A, S3C; boxes). For example, the first neuron in Figure 2A (top) fired transiently both when the participant heard “*earth’s the matter*” (green box) and when they spoke the same phrase (red box). Aligning the timings of individual phonemes between the heard and the spoken phrases (Figure 2B, top; Methods) reveals that the listening response lags “*the*” in “*earth’s the matter*”, while the speaking response occurs before/during “*the*”. The second neuron (Figures 2A,B; bottom) showed similar pairs of listening and speaking activity but at “*will*” and at “*me*”, respectively, in the sentence “*will you tell me why*”. The causality in listening-lag and speaking-lead activity to the same speech elements suggests the dual (auditory and motor) nature of these mirror neurons. While mirror neurons have been observed in other behavioral domains and brain areas, direct evidence for their existence for human speech has not been shown previously.

**Figure 2.**
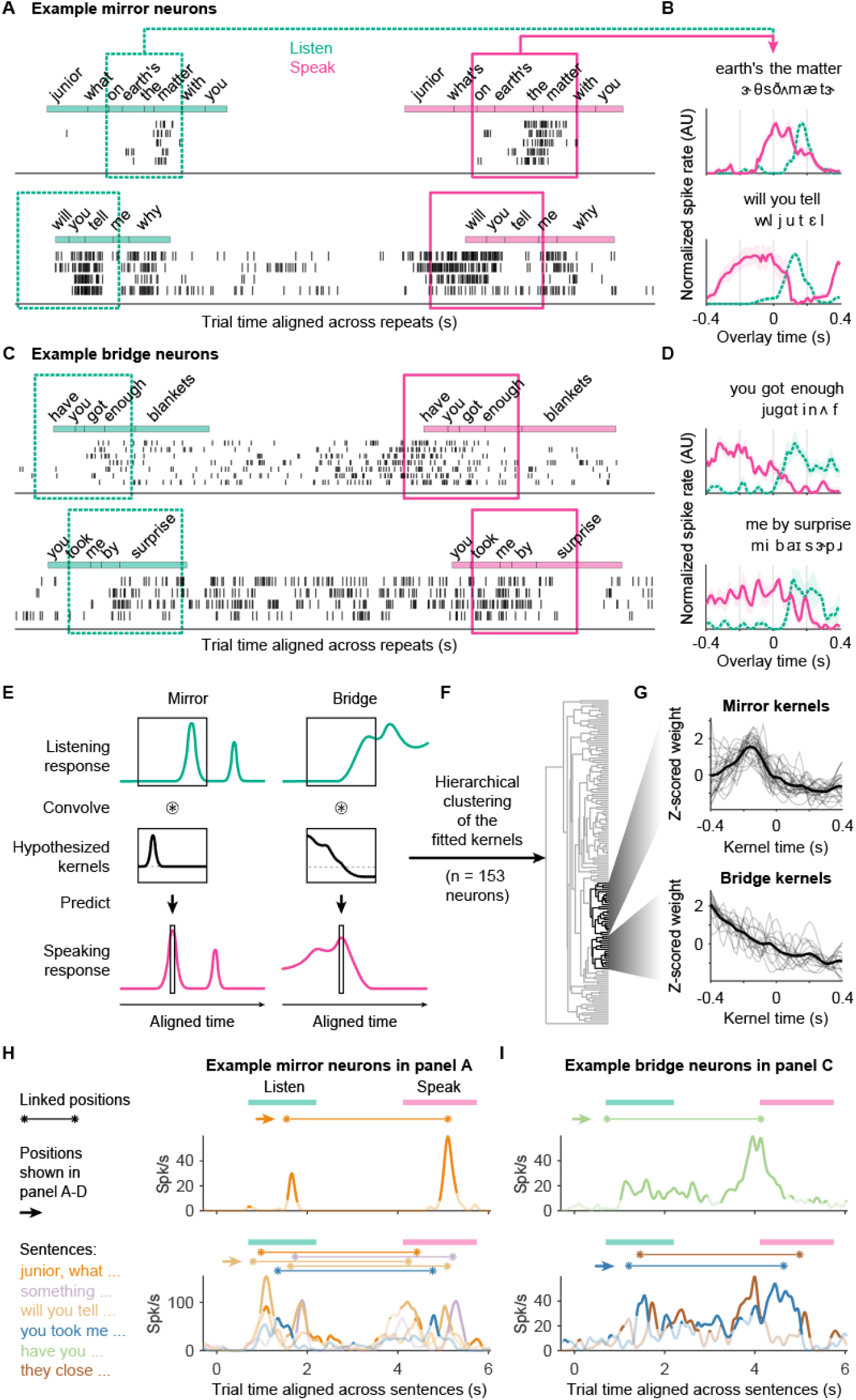
“Mirror” and “bridge” neurons link speech elements from perception to production. (A) Spike rasters of two example mirror neurons each firing to similar parts of the sentences during listening and speaking. (B) Zoomed in overlays of listening (dashed green) and speaking (solid magenta) responses (mean ± SD) for the neurons in (A), showing that the speaking activity precedes the listening responses. (C,D) Similar to (A,B) but showing two example bridge neurons with elevated activity between the same speech elements in listening and speaking, across delay and initiation. (E) Schematics of the modeling procedures and the hypothesized kernels for mirror (left) and bridge (right) neurons. (F) Hierarchical clustering on the kernels of listening-and-speaking responsive neurons produces clusters of mirror (upper) and bridge (lower) kernels. (G) Overlays of mirror (top; n = 25) and bridge (bottom = 16) kernels from their clusters in (F). Thick traces indicate the mean. (H) Trial-averaged responses of the two example mirror neurons in (A,B) to sentences (traces) that have linked activity (connected asterisks). Darker segments of the traces indicate responses that are sentence-specific (outside 95% bootstrap confidence interval of trial-shuffled responses). Sentences without linked activity are omitted to avoid clutter. (I) Same convention as in (H) but showing responses of the two example bridge neurons in (C,D).

More surprisingly, another group which we called “bridge” neurons maintained elevated firing between the same part of the sentence in listening and speaking, bridging over delay and initiation phases of the task (Figures 2C,D, S3D). The first example neuron (Figures 2C,D; top) started firing after the participant heard “*you*” in the phrase “*have you got*”. The elevated spiking remained through delay and initiation, and decreased when the same speech sound was produced during the speaking phase. The second neuron (Figures 2C,D; bottom) showed similar bridging activity between listening and speaking but for “*by*” in “*me by surprise*” of a different sentence. The presence of mirror neurons suggests there may be shared representations during the perception and production of the same speech content, while bridge neurons may contribute in parallel to carrying these elements from the moment of perception to the time of production.

To objectively identify all candidate mirror and bridge neurons, we used an unbiased approach by modeling the relationship of each neuron’s activity between perception and production (Methods; Figure 2E). For each unit, perception responses in a moving window centered at a given sentence position is used to predict a single value of production activity at the same position via a linear kernel. A mirror neuron would result in a narrow positive bell-shaped kernel, offset to the past (left) of the current sentence position. Such a kernel translates a trailing perception response into a preceding production activity. A bridge neuron, however, would result in a kernel with downward slope from positive to negative weights. This approximates the transformation from a prolonged response after perceiving given elements and before producing the same elements. This model is not preconfigured to look for neurons with mirror, bridge, or other temporal profiles, but attempts to identify any linear relationship between a response to speech content at a particular sentence position during listening and a response to the same speech content during speaking.

These models were fitted for all neurons that were responsive during both listening and speaking. To identify all possible linker types in an unsupervised manner, we performed hierarchical clustering across all fitted kernels (Figure 2F). Three interpretable clusters were identified: mirror type kernels (Figure 2G, top; kernel peak time at −0.14 ± 0.11 s, mean ± SD), bridge type kernels (Figure 2G, bottom), and auditory kernels (see next section). The kernels outside the three clusters consisted of noisy, acausal, or overfit weights (Figure S4B). These kernels were uninterpretable and may have resulted from 1) limited statistical power with certain neurons, 2) a lack of relationship between listening and speaking responses, or 3) the relationship cannot be approximated using a linear kernel.

The successfully fitted kernels allowed us to systematically identify the locations of speech sounds that are linked between listening and speaking (Methods). The first example mirror neuron (Figure 2H; top) has only one pair of linked positions in one sentence (“*junior, what …*”), whereas the second mirror neuron (Figure 2H; bottom) shows linked positions in four different sentences with one sentence (“*will you …*”) having two separate pairs of links. To directly evaluate the neuronal coding of speech information across task phases, we used mirror neuron activity to decode the sentence identity at the single-trial level (Methods). We found that decoders fitted during listening can both decode sentence identity from the held-out trials during listening but also during speaking, and vice versa (Figure S3E; left). This decoder generalization demonstrates that mirror neurons used a stable code to represent speech content between listening and speaking (Figure S3F; top).

The locations of linked speech sounds were also identified for bridge neurons. The first bridge neuron (Figure 2I; top) links a pair of positions in only one sentence (“*have you…*”), whereas the second bridge neuron (Figure 2I; bottom) links in two different sentences. We performed the same sentence identity decoding and found pairwise generalization among listening, delay, initiation, and speaking (Figure S3E; right). This shows that bridge neurons used a largely stable code to carry speech information from listening to speaking (Figure S3F; bottom), which can support the maintenance of verbal working memory necessary for performing this delayed sentence repetition task.

These results demonstrate the linking of a diverse set of speech elements between listening and speaking. The presence of speech mirror and bridge neurons reveals shared and linked representations of speech content between sensory input and motor output.

### Auditory neurons respond to both external and feedback of self-produced speech sounds

In contrast to the mirror and bridge neurons that fired preceding speech production, we found auditory neurons responding with near identical lags to both external and self-produced speech sounds during listening and speaking, respectively (Figure 3A,B, S3A). The example neuron (Figure 3A,B) showed auditory responses to the sound /s/ in “*this house*” and in “*to search*”. The time courses of listening and speaking responses overlap after aligning the exact timings of the external and self-produced speech at single-phoneme precision (Figure 3B).

**Figure 3.**
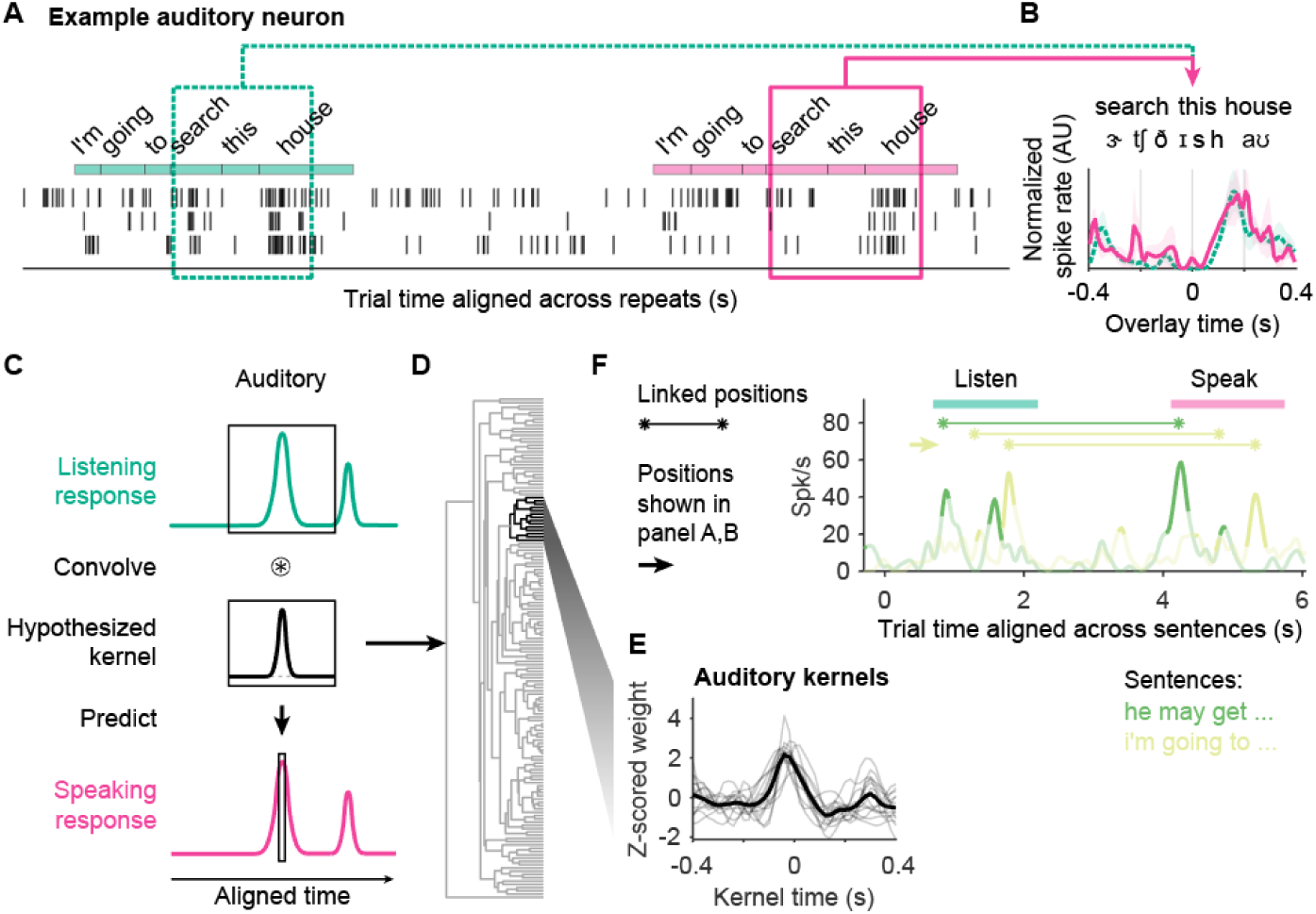
Auditory neurons respond to both external and self-produced speech sounds. (A) Spike rasters of an example auditory neuron firing to the same speech sounds during listening and speaking. (B) Zoomed in overlays of listening (dashed green) and speaking (solid magenta) responses (mean ± SD) for the neuron in (A), showing that the listening and speaking responses well overlap. (C) Schematics of the modeling showing the hypothesized kernel for auditory neurons. (D) The same hierarchical clustering from Figure 2F also produces an auditory cluster. (E) Overlay of auditory kernels (n = 14) from the cluster in (D). Thick trace indicates the mean. (F) Trial-averaged responses of the example auditory neuron in (A) to sentences (traces) that have linked activity (connected asterisks). Darker segments of the traces indicate responses that are sentence-specific (outside 95% bootstrap confidence interval of trial-shuffled responses). Sentences without linked activity are omitted to avoid clutter.

The transformation kernel that captures such an auditory neuron seems similar to that of a mirror neuron but the bell-curve would be centered around zero time offset (Figure 3C) instead of being shifted to the past as in mirror kernels (Figure 2E; left). The same hierarchical clustering described in the previous section also identified a cluster of auditory kernels (Figure 3D,E). We compared the peak times between auditory kernels (−0.01 ± 0.09 s; mean ± SD) and mirror kernels (−0.14 ± 0.11 s; mean ± SD) and found a significant difference (p = 2.3e-5, two-tailed Wilcoxon rank-sum test). That is, these auditory neurons are firing in response to speech sounds (i.e. auditory feedback), and do not have a clear role in feedforward speech production like mirror neurons. Auditory responses to the same sounds during listening and speaking can occur in multiple sentences and more than one part of a sentence (Figure 3F).

### Distribution of mirror, bridge, and auditory neurons across cortical regions and cortical depth

We next studied whether these different functional types are enriched in specific regions or cortical depths. We found that bridge neurons were exclusively located in PrCG, with 87.5% located in the mPrCG (Figure 4A). Similarly, 76.0% of mirror neurons were in PrCG, with 89.5% of those in mPrCG (Figure 4B). Although the posterior IFG was originally theorized to contain cells with mirror properties and to mediate phonological mapping ^16^, we found no neurons in IFG-op that met the criteria for mirror or bridge neurons (Figure 4D). These results suggest a distinctive role for the mPrCG, as it is the predominant region to both represent perceived and produced speech content in a shared coding space and show persistent activity that can support working memory of speech.

**Figure 4.**
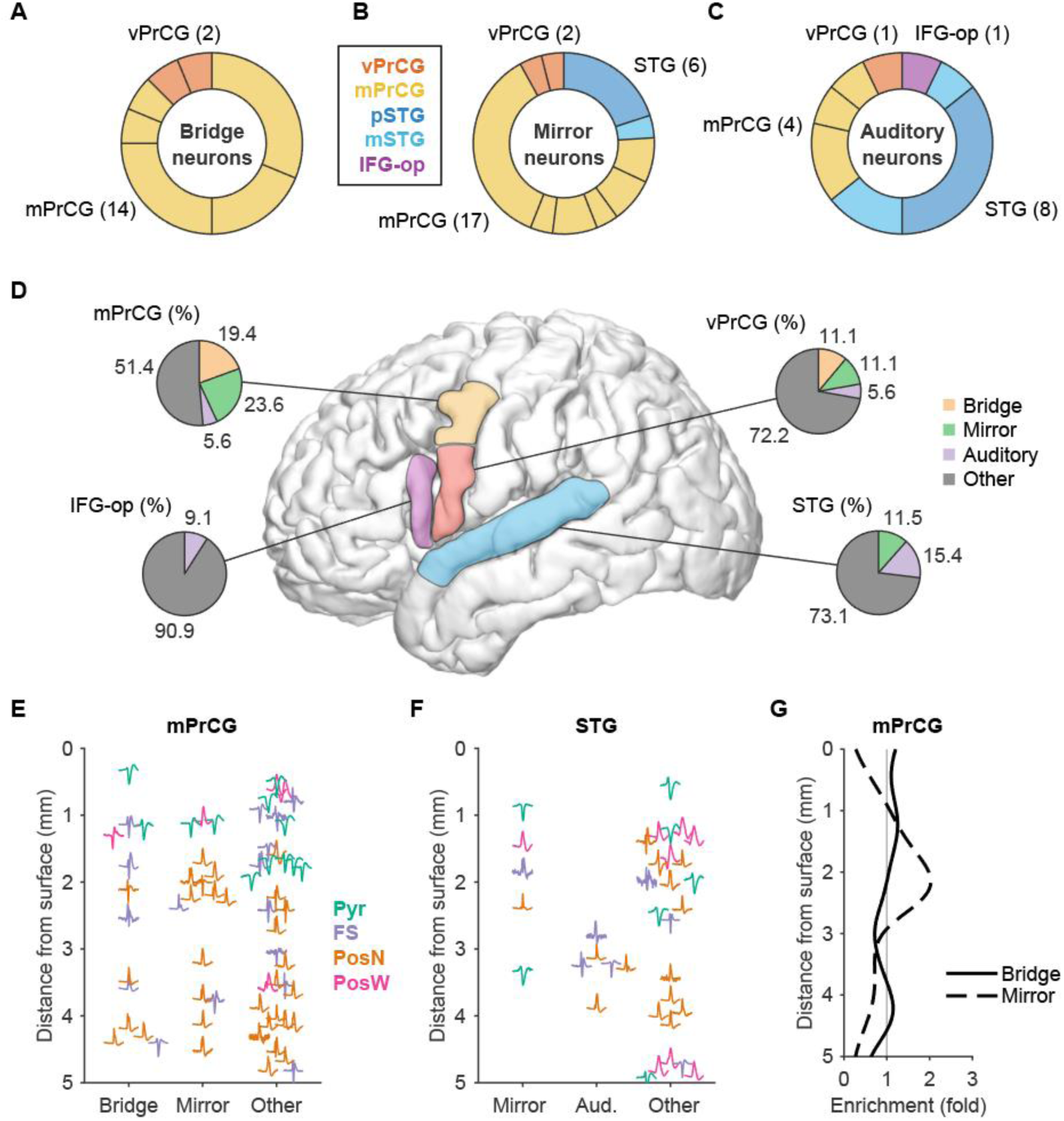
Distribution of mirror, bridge, and auditory neurons across cortical regions and depth. (A) Cortical origin of identified bridge neurons. Colors indicate different cortical regions. Each section is a recording. The number of neurons from each region is indicated in the parentheses. (B,C) Similar to (A) but for mirror and auditory neurons, respectively. (D) Percentages of mirror, bridge, and auditory neurons out of all listening-and-speaking responsive neurons in each of the four regions. (E) Peak normalized median spike waveforms of bridge, mirror, and other listening-speaking responsive units in mPrCG at the recorded cortical depths. Colors indicate different waveform types. (F) Similar to (E) but for mirror and auditory units in STG. (G) Distributions of bridge (solid; n.s.) and mirror (dashed; n.s., p = 0.08) neurons across the cortical depth of mPrCG, relative to all task-response neurons. Distribution of auditory neurons is not estimated due to a low sample count (n = 4 units). Two-sample K-S test.

Indeed, neurons in the auditory cluster were predominately in the STG (including both middle and posterior sites; a small number were also in the PrCG) (Figure 4C). Further consistent with the role of the pSTG in sensorimotor functions, we found a small number of mirror neurons in this area (Figures 4B,D).

Bridge and mirror were found in a broad range of cortical depth and with all four spike waveform types (Pyr, FS, PosN, and PosW in Figures 4E,F). However, we did not find mirror or bridge neurons concentrated in particular cortical depths. Specifically, we fitted the depth distributions for each functional group and normalized them by the distribution for all task-responsive units. In mPrCG (Figure 4G), mirror neurons show a trend, though not significant (p = 0.08), of being concentrated in the middle cortical layers (around 2 mm from cortical surface). The distribution of bridge neurons did not differ from the distribution of overall task responsive neurons. In STG, we did not have enough samples to quantify the cortical depth distributions, though all the auditory neurons were located in deeper layers. Thus, in general cell types and cortical depth do not seem to be defining features of mirror, bridge, and auditory neurons.

### Latent population dynamics define task phases and sustained memory activity

Most of the analyses so far have focused on the activity of single neurons. Recent studies in delayed reaching tasks have suggested that neuronal population activity can reveal important latent factors that reflect emergent computational properties above what can be deduced from single neuron analyses ^17^. Because many neurons are active during both the delay period and during movement ^18^, with no clear similarity between these two activities, individual-neuron responses can be difficult to interpret. Like the delayed reaching task, our delayed repetition task was designed to explore processes related to listening and speaking execution, but also the internal processes of working memory, preparation, and initiation. To identify distinct latent factors underlying these processes, we used Sparse Component Analysis (SCA) ^19^ which is a unsupervised dimensionality reduction method that encourages partitioning of population activity into temporally distinct components across the recordings.

Using SCA, we reduced neuronal population activity separately in each of the four cortical regions to 12 components (Methods). The latent factors in the mPrCG population fall into two distinct categories. The first category includes four phasic components (Figure 5A; ranked by descending variance explained), each primarily showing activations to a single, specific speech processing phase. The first two components show activation during speaking, highlighting that the largest explained variance across mPrCG neurons is for speech production. The third and fourth components are active during listening and initiation, respectively. These latent components at distinct task phases are consistent with the temporal motifs of single neuron responses in mPrCG (Figure 1C), and demonstrate that these functions are generated from intermixed neuronal populations. vPrCG has similar phasic components as mPrCG (Figure S5C) except for reduced activations in the listening phase (p = 7e-28, D* = 0.77, two-sample two-tailed KS test, n = 112 single-trial responses projected to mPrCG component #3 and n = 112 to vPrCG component #6). IFG-op has only one phasic component that distinguishes speaking and the combined phases outside speaking (Figure S5D), consistent with the clustering result of neuronal responses shown in Figure 1E. STG shows phasic components in listening and speaking (Figure S5E).

**Figure 5.**
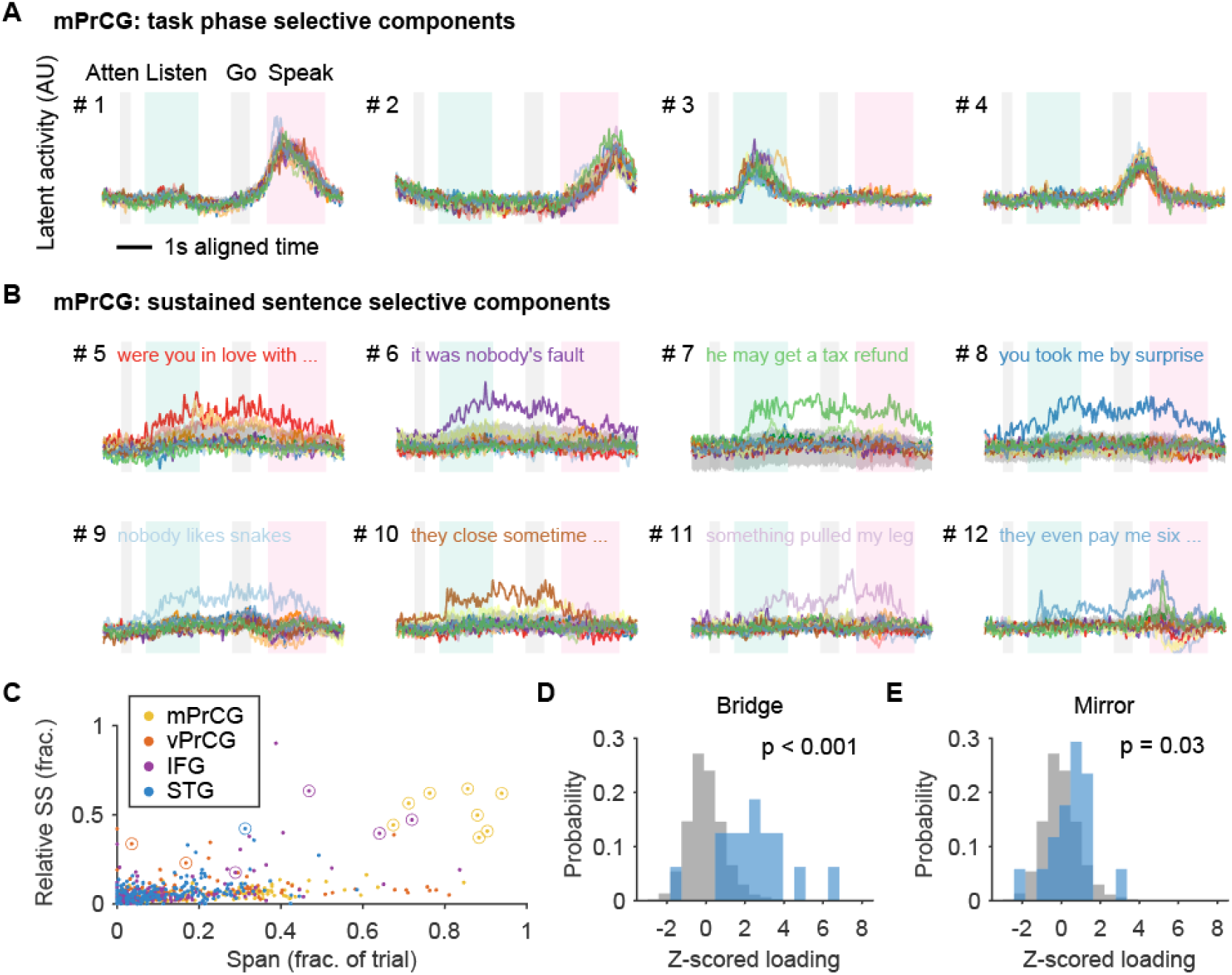
Population activity in mPrCG contain sustained sentence-selective dynamics. (A) Latent activity of SCA components #1-4 from mPrCG populations. Colored traces represent the 14 different sentences. Shading around traces represents the 95% bootstrap confidence interval of projected activity using shuffled trials. In each plot, the shaded periods from left to right indicate attention cue (Atten), listening (Lisen), go cue (Go), and speaking (Speak). (B) Same conventions as in (A) but showing components #5-12. In each plot, the text labels the sentence with significant high activation across task phases from listening to speaking. (C) Each dot represents the latent activation for a unique sentence in a component. y-axis, the fraction of sum-of-squared (SS) of an activation out of the total SS across all sentences in a given component; x-axis, the span of the activation during a trial. Circled activations are those with significant high activation across task phases, such as those labeled in (B) by texts. (D) Loadings of bridge neurons (blue) in components #5-12 for the preferred sentences (n = 16 loadings) compared with that of all neurons (gray; n = 4000 loadings). Two-sample K-S test. (E) Similar to (D) but comparing loadings of mirror neurons (blue; n = 17 loadings).

The second category (i.e. the remaining eight) of mPrCG components (Figure 5B) each shows activation to a small number, usually just one sentence. Unlike the task phase-specific temporal profiles above, these components are sentence-specific and sustained. The components ramp up during listening, persist over the memory periods, and ramp back down towards the end of speaking. Though vPrCG, IFG-op, and STG all have latent activations that are selective to specific phases (Figures S5C-E), only mPrCG exhibits prevalent sustained sentence-selective activations (Figures 5C). Activations in the other three regions (Figure 5C) are usually either not selective to specific sentences (low fraction of single-sentence sum-of-squared activity out of all sentences) or sentence selective but only within specific task phases (low span in the period from perception onset to production offset).

To better understand the contribution of bridge and mirror neurons to the sustained sentence-selective components, we examined the loadings (measure of contribution) of each neuron to these components. We found that bridge neurons provided significantly higher loadings for sentence-selective components than randomly selected neurons (2.3 ± 1.9, mean ± SD, p < 0.001; Figure 5D). Whereas the mirror neurons, which do not have persistent activity, had small but statistically significant contributions (0.5 ± 1.1, mean ± SD, p = 0.03; Figure 5E). Given the strong contribution from bridge neurons, we wondered whether the presence of these components is solely driven by the small number of bridge neurons that maintained speech working memory (Figure S3E,F), or additionally reflects dynamic working memory code ^20^ supported by a broader range of cells. To test this, we performed SCA on mPrCG population excluding all the bridge neurons (Figure S5F). These components are largely preserved but with minor degradation (median 9% (IQR 5-54%) reduction in latent activation; p = 0.008, two-tailed Wilcoxon signed rank test, n = 8 activations). These results demonstrate that mPrCG can support speech working memory with both persistent activity from single bridge neurons and a dynamic population code.

### Speech feature selectivity in precentral gyrus neurons

Previous work using electrocorticography showed tuning to acoustic spectral features, like vocal pitch during listening and speaking ^7,21^, and somatotopic representations of articulators centered around the central sulcus (primary motor cortex) during speaking ^2,3,22^. In addition to these candidate representations, PrCG neurons may encode a representation that corresponds to sequences at the segmental level (phoneme or syllabic units), proposed to be relevant for speech planning ^23^.

Our task was designed to study neuronal activation across listening and speaking, using 14 sentences repeated over multiple trials. We did not have enough time during recordings to comprehensively sample a large speech feature space. Nonetheless, given these constraints, we explored to what extent single PrCG neurons encode auditory spectral cues, vocal tract articulatory movements, or phoneme features.

To address this, we fitted separate encoding models for each neuron using features either from the Mel spectrogram, the estimated articulatory kinematic trajectories (AKTs) and their rate of change, or single-phoneme features (Figure S6A; Table S2). Neurons tuned to these quickly varying features should exhibit time-locked responses. Therefore, there must exist an optimal time offset between features and responses where the goodness-of-fit (*r*^2^) is maximized and statistically significant (e.g. Figure S6B; Methods). Significant encoding is determined by comparing model fits to shuffled null distribution. All responsive neurons are included given each task phase of interest.

In the PrCG recording sites, 4.5% of the neurons have significant encoding to AKT features (Figure 6A, top left). Across PrCG neurons, model weights showed heterogeneous tunings to different combinations of articulatory features (Figure S6C). As an example (Figure S6D), the spiking of a vPrCG neuron was positively tuned to the size of jaw opening (Figure S6E, top) and negatively tuned to tongue tip curvature (Figure S6E, bottom). During listening, only 1.4% (or n = 5) of the listening responsive PrCG neurons show significant encoding of spectral features (Figure 6A, bottom left; Figures S6F-H). In contrast, 20% of STG neurons exhibited spectral tuning (Figure 6A, bottom right).

**Figure 6.**
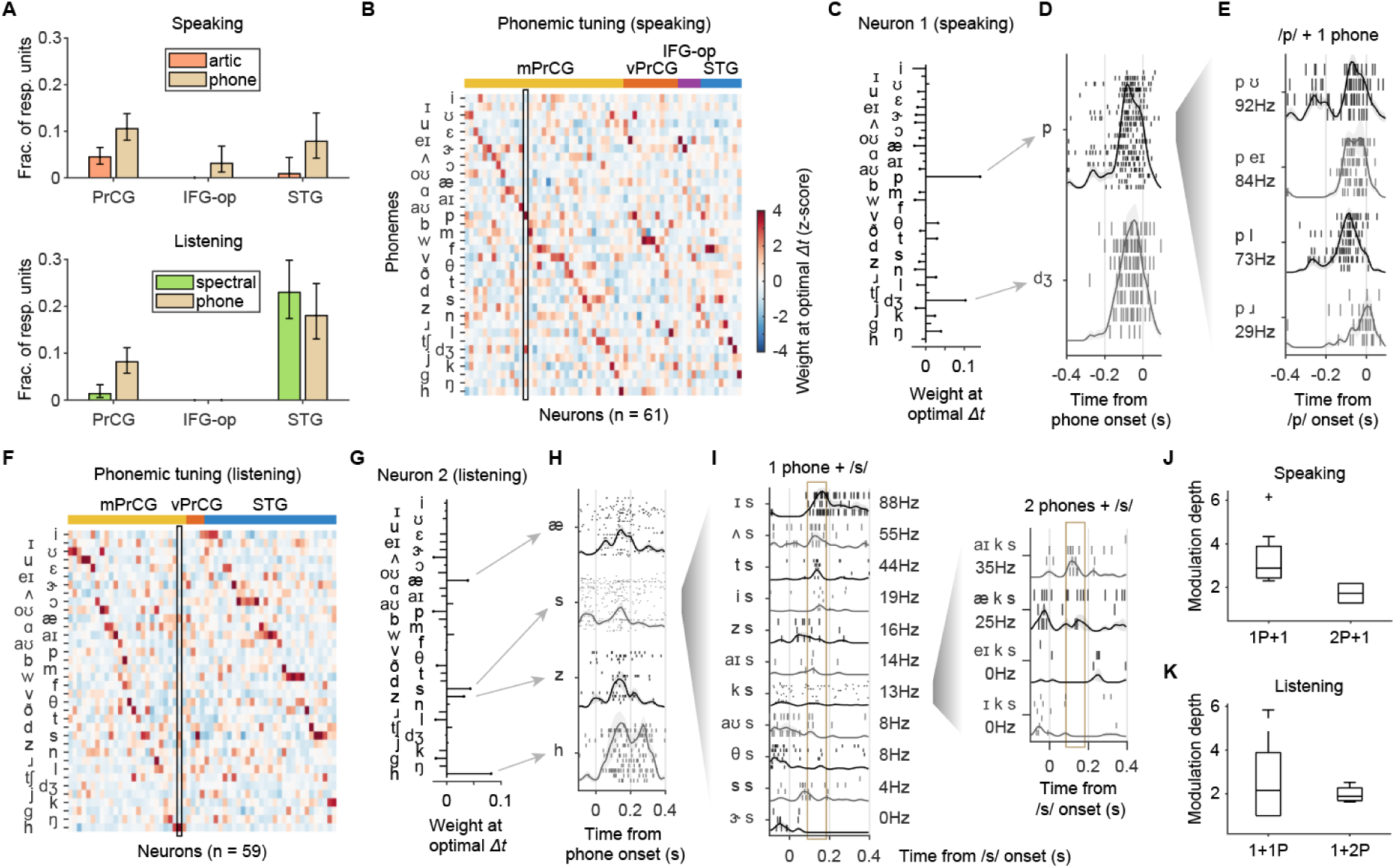
Speech feature selectivity in precentral gyrus neurons. (A) Fractions of responsive units that show time-locked spectral, articulatory, or phonemic encoding for speech production (top) or stimulus perception (bottom). (B) Normalized model weights across neurons (columns, groups by regions and sorted by peak weight position) that have significant phonemic encoding for speaking. The neuron highlighted in the box is expanded in (C-E). (C) Weights of the phonemic model highlighted in (B). Weights with stem heads are statistically significant. (D) Spike rasters and PETHs (mean ± SEM) of the two best tuned phonemes in (C). (E) Spike rasters and PETHs (mean ± SEM) showing responses at /p/, broken down by different 2-phone sequences. Labeled spike rates are measured at optimal *Δt* (Methods). (F-H) Similar to (B-D) but for phonemic models fitted during listening. (I) Spike rasters and PETHs (mean ± SEM) showing responses to unique 2-phone sequences that end with /s/ (left), and 3-phone sequences that end with /k s/ (right). Vertical boxes mark around the optimal *Δt*. Same conventions as (E). (J) Relative modulation depth during speaking when branching from single phonemes (left) or 2-phone sequences (right) to include one extra phoneme from the context. N = 11 units. (K) Similar to (J) but for listening. n = 12 units.

We next fitted models with phonemes, which are the segmental units of consonants and vowels. Higher proportions of PrCG neurons showed phonemic encoding during listening (8.2%) or speaking (10.5%) (Figure 6A). At the population level, PrCG neurons showed diverse tuning patterns with top weights covering the majority of phonemes (Figures S6B,F). For example, a speech production neuron in mPrCG (Figures 6C,D) was selective to two phonetically very dissimilar phonemes, /p/ and /dʒ/ (i.e. “j” sound in “junior”). A speech perception neuron in mPrCG (Figure 6G,H) was selective to three consonants (/s/, /z/, /h/) and a vowel (/æ/). These combinations in phonemic tuning cannot be fully explained by a consistent spectral or articulatory feature.

Furthermore, we found that responses of PrCG neurons to their tuned phonemes can be modulated by surrounding phonetic context. For the example speech production neuron shown above (Figure 6C), its time-locked responses to /p/ can vary strongly from 29 to 92 spikes/s (p = 0.03, Kruskall-wallis test; Figure 6E) depending on different upcoming phonemes (i.e. /p/ + /ʊ/, /eɪ/, /l/, or /ɹ/). The example speech perception neuron (Figure 6G) showed a wide range of modulations at the levels of both 1-phoneme + /s/ (Figure 6I, left) and 2-phoneme + /s/ (Figure 6I, right).

We quantified the extent of response modulation from single phonemes to 2-phoneme (2P) sequences and from 2P to 3P sequences for each neuron that had significant phonemic encoding (Figure 6J,K; Methods), and observed strong 2P (2.2 (1.0-3.9) folds change in spike rates for listening and 2.7 (2.3-3.7) for speaking; median (IQR)) and 3P (1.9 (1.7-2.3) folds for listening and 1.7 (1.3-2.2) for speaking; median (IQR)) modulations. Overall, these findings demonstrate selectivity in the precentral gyrus neurons to speech features at the segmental level and the effects of sequence modulation.

## Discussion

In summary, our large-scale neuronal recordings reveal the cellular basis of a speech circuit that tightly links sensory input with motor output. This circuit leverages an internal map of phonological elements, enabling seamless integration between speech perception and production. The discovery of mirror and bridge neurons within the middle precentral gyrus (mPrCG), alongside persistent and content-specific population dynamics, highlights the mPrCG’s crucial role in both real-time speech processing and working memory.

These findings contribute significant updates to established models of speech production, imitation, and vocal learning. Notably, Levelt and colleagues proposed a process of “phonetic encoding,” where phonological representations are mapped onto motor commands via a “mental syllabary” ^24,25^. Similarly, the DIVA model posits that a “speech sound map” drives production, with cells that may function as mirror neurons, responding to the same sounds during both listening and speaking ^16^.

While mirror neurons have been extensively characterized in animal models ^26–29^, their existence in humans remains controversial, with limited evidence beyond grasping and facial movements ^30^. Mirror-like mechanisms for language have been postulated in Broca’s area ^31–34^. However, our data did not reveal mirror neurons in Broca’s area. Broca’s area has also been proposed to mediate phonological processing and working memory that are important for speech planning and articulation. However, the absence of bridge neurons and the minimal persistent population dynamics in this area does not support these roles. This aligns with accumulating evidence that Broca’s area injury or perturbation does not necessarily lead to speech disfluency ^35–37^. Instead, we found speech-related mirror neurons predominantly in the PrCG and posterior superior temporal gyrus (pSTG), both of which are connected via white matter tracts ^38^ and implicated in speech production deficits.

Speech repetition and imitation rely on maintaining auditory information in working memory until it can be articulated. Bridge neurons, with their persistent and content-specific activity, may facilitate this process. Unlike mirror neurons, bridge neurons were found exclusively in the PrCG (especially in mPrCG), where they supported sustained representation across all task phases: listening, delay, initiation, and speaking. The mPrCG’s unique role in linking sensory, motor, and cognitive representations of speech is further emphasized by its content-specific population dynamics. Working memory in the mPrCG appears to rely on both persistent single-neuron activity and dynamic population-level coding through latent components, consistent with theories of working memory encoding ^20^.

Clinically, the PrCG has traditionally been considered part of the primary motor cortex ^39,40^, but only its posterior bank exhibits primary motor cortex characteristics ^41^. Our findings align more closely with premotor functions ^42–44^. Notably, more neurons in the PrCG were tuned to complex speech sequences rather than simple articulatory movements or acoustic features. In contrast, previous studies have shown that the coding of basic articulatory and acoustic features was more prevalent in the posterior PrCG and adjacent postcentral gyrus ^3,7,45^. The relative scarcity of articulatory-tuned neurons in our dataset likely reflects the anterior positioning of our recording sites, as posterior regions were preserved to avoid motor impairments during surgery ^46^.

The speech sequence-dependent activity of PrCG neurons is consistent with the premotor role of PrCG in higher-order speech planning and sequencing, as seen in cases of pure apraxia of speech following PrCG resection ^47–51^, where patients struggle with planning and sequencing speech despite intact orofacial strength and language functions. Given the limited recording time and specific task design, we did not have the opportunity to further explore alternative mechanisms, such as hyper- or hypo-articulation in certain words ^52^, or variations in syllabic stress patterns ^53^, that may underlie the phonemic sequence-dependent activity. Future studies incorporating richer and/or more controlled stimulus sets will be essential for a more comprehensive understanding of neuronal representations within the PrCG and the broader speech network.

Overall, our study offers a novel framework for understanding the complex neural mechanisms underlying speech. By revealing how perception, production, and memory are integrated at the cellular level, these findings open new avenues for exploring the intricate neural architecture that supports human communication.

## Acknowledgements

We thank all the participants who volunteered in this study, Adib Abla for collaboration on participant recruitment and surgical operations, Josh Glasser for discussion on the Sparse Component Analysis, Ueli Rutishauser for discussion on working memory analysis, Jessie Liu for extensive suggestions on writing the manuscript, Cathryn Cadwell for assistance with the histology pipeline, and Keith Johnson for advice on phonetic analysis. This work was supported by grants from Prometheus (Fred Ehrsam), the Chan Zuckerberg Biohub, and Weill Neurohub.

## Author Contributions

D.X. and E.F.C. conceived the experiments. E.F.C., J.E.C., M.K.A., and S.L.H. performed the surgeries. D.X., J.E.C, E.F.C., Q.R.G, and others collected the data. D.X., S.L.M., and Q.R.G. developed the software and preprocessed/curated the data. D.X. analyzed the data with help from S.L.M., A.B.S, Y.Z., and M.K.L.. D.X. visualized the results. D.X., E.F.C., Q.R.G and Y.Z. wrote the paper which was revised by all authors. E.F.C. supervised the project.

## Declaration of Interests

E.F.C. is a co-founder of Echo Neurotechnologies Corporation, LLC. The remaining authors declare no competing interests.

---

**Figure S1.**
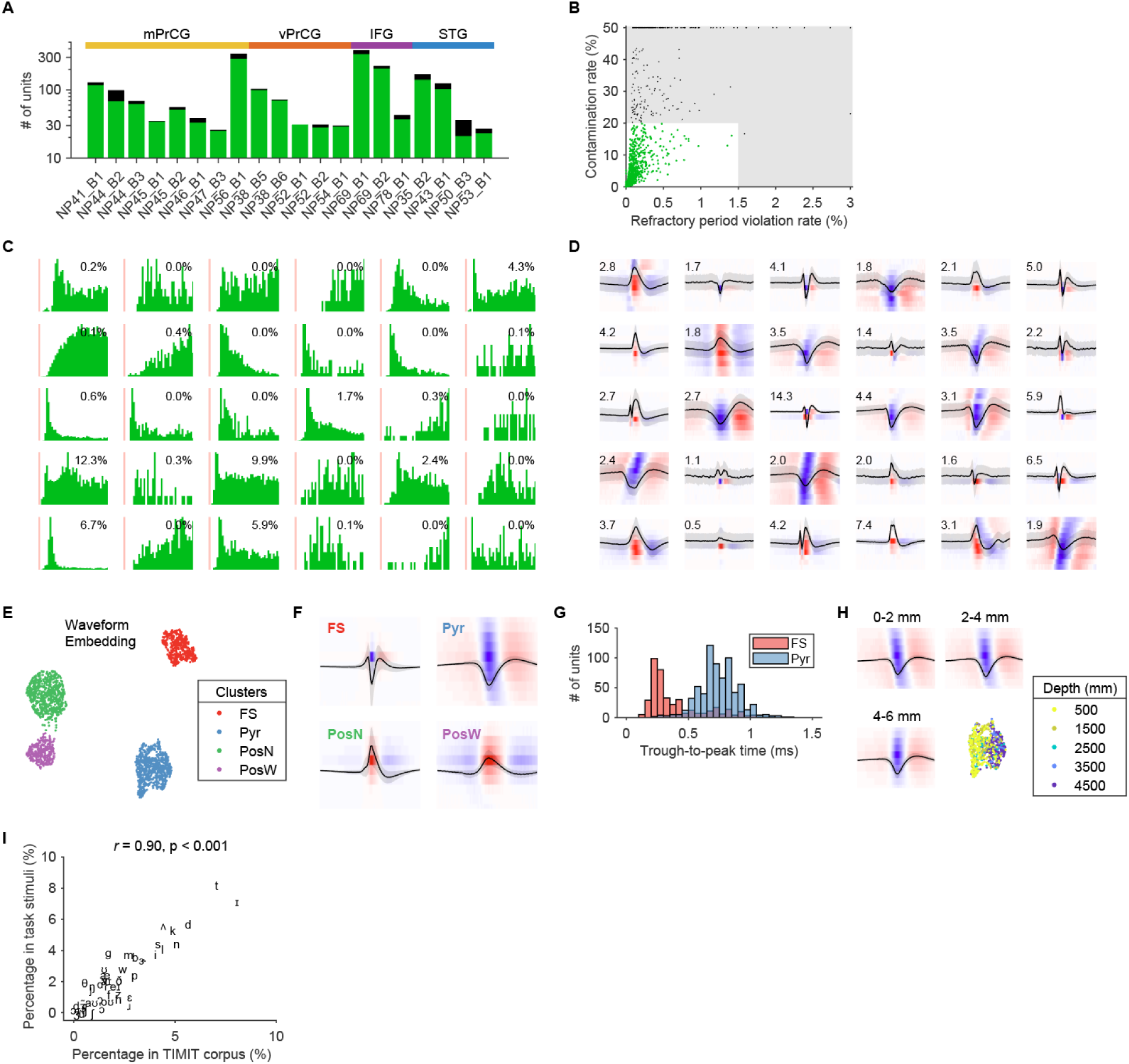
Recording yield and unit properties. (A) Yield of single-(green) and multi-units (black) for each recording included in this study. (B) Scatter plot showing refractory period violation rate (RPV) and contamination rate (CR). Green dots represent single-units, black multi-units. Suprathreshold ranges are shaded. (C) ISI histograms of randomly selected units. Red shades indicate 1.5 ms refractory periods. Percentages indicate the contamination rates. (D) Median spatiotemporal spike waveforms of the same units in (C). Numbers indicate the waveform SNR. (E) UMAP embedding of spatiotemporal spike waveforms followed by Gaussian mixture model clustering results in four well-isolated clusters (colored). FS, putative fast spiking neuron; Pyr, putative pyramidal neuron; PosN, narrow positive waveform unit; PosW, wide positive waveform unit. (F) The average waveforms of the four clusters. Heatmaps show the normalized spatiotemporal waveforms. Black traces show the waveforms (mean ± SD) from the center electrodes. (G) Units in FS cluster and Pyr cluster show well separated spike trough-to-peak time. (H) Spatiotemporal waveforms of Pyr vary as a function of cortical depth. Average waveforms of different depth ranges are shown using the same conventions as in (F). Lower right shows the Pyr waveform cluster in (E) but colored by cortical depth. (I) Scatter plot comparing the percentage of each unique phoneme out of all phonemes in the full TIMIT corpus versus that in the selected sentences of the present task. Pearson’s correlation, n = 43 unique phonemes.

**Figure S2.**
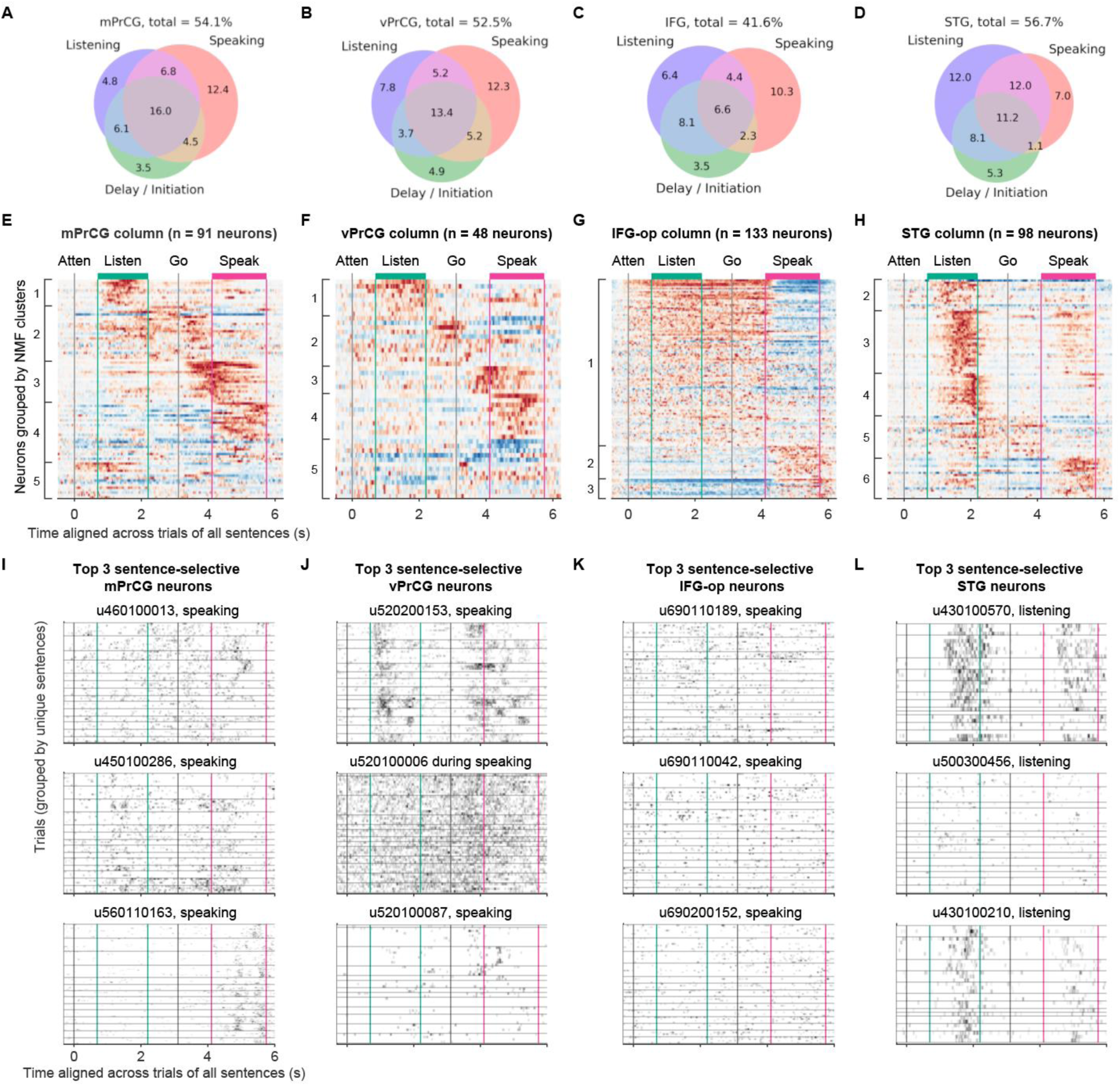
Neuronal responses across task phases. (A) The percentages of neurons out of all recorded mPrCG neurons that are responsive to different task phases. (B-D) Similar to (A) but for vPrCG, IFG-op, and STG, respectively. (E) Average evoked activity of task responsive mPrCG neurons from a single cortical column, grouped by NMF cluster membership and sorted by decreasing loadings. (F-H) Similar to (E) but for vPrCG, IFG-op, and STG, respectively. (I) Three mPrCG neurons with the highest significant chi-square statistic of the Kruskal-Wallis test for sentence-selectivity. Each heatmap shows single-trial spike rates of a neuron. Trials are grouped by the same sentences. The task phases where the highest chi-square statistic were found are indicated in subtitles. (J-L) Similar to (I) but for vPrCG, IFG-op, and STG neurons.

**Figure S3.**
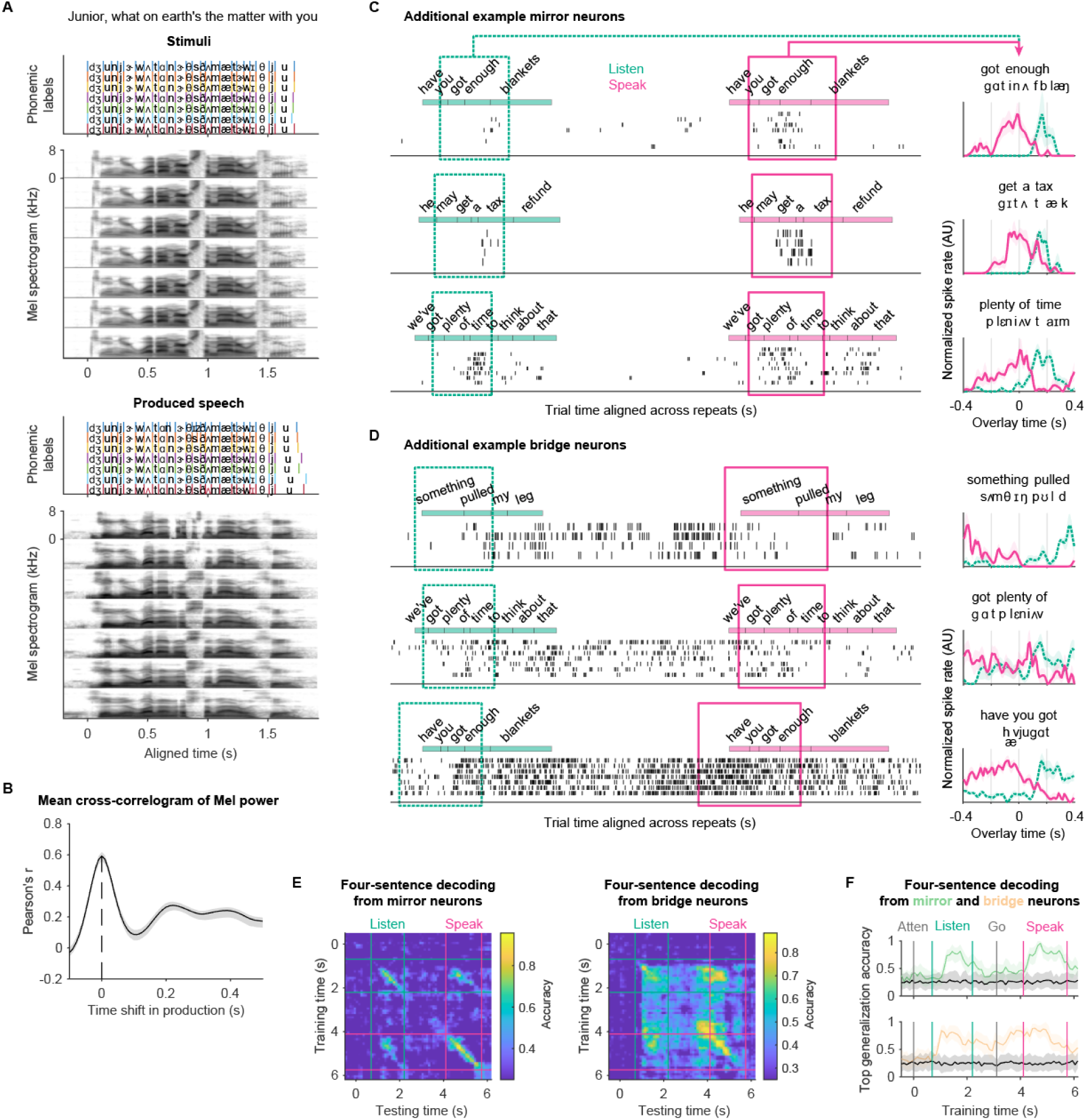
Additional examples of mirror and bridge neurons. (A) Individual trials of produced speech (bottom) are well aligned with speech during listening (top). (B) Cross-correlogram (mean ± 95% confidence interval; n = 20 recordings) between the Mel spectrogram power of stimuli and produced speech shows a peak at zero second (dashed line), indicating unbiased alignment. (C) Three additional examples of mirror neurons. Same conventions as Figures 2A,B. (D) Three additional examples of bridge neurons. Same conventions as Figures 2C,D. (E) Left, four-sentence cross-time decoding accuracy using the 25 mirror neurons. Each value in the heatmap is the accuracy of a decoder trained at one task time (y-axis) and tested at the same or another task time (x-axis). Right, similar to Left but using the 17 bridge neurons. (F) Best generalized four-sentence decoding accuracy for decoders trained at each time point (x-axis) with mirror (green) or bridge (yellow) neuron responses.

**Figure S4.**
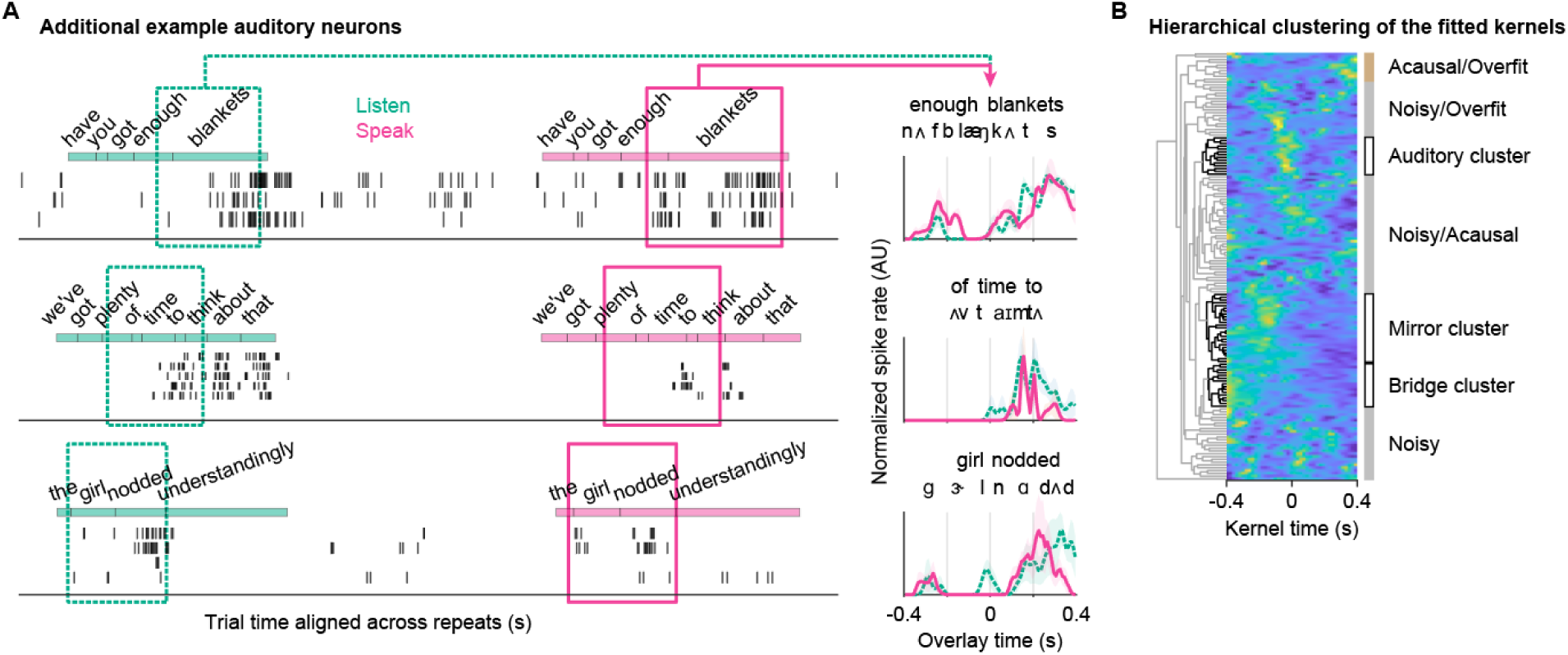
Additional examples of auditory neurons. (A) Three additional examples of auditory neurons. Same conventions as Figure 3A,B. (B) Timeseries of all the 153 hierarchically clustered kernels. Uninterpretable kernel groups are labeled alongside the bridge, mirror, and auditory clusters.

**Figure S5.**
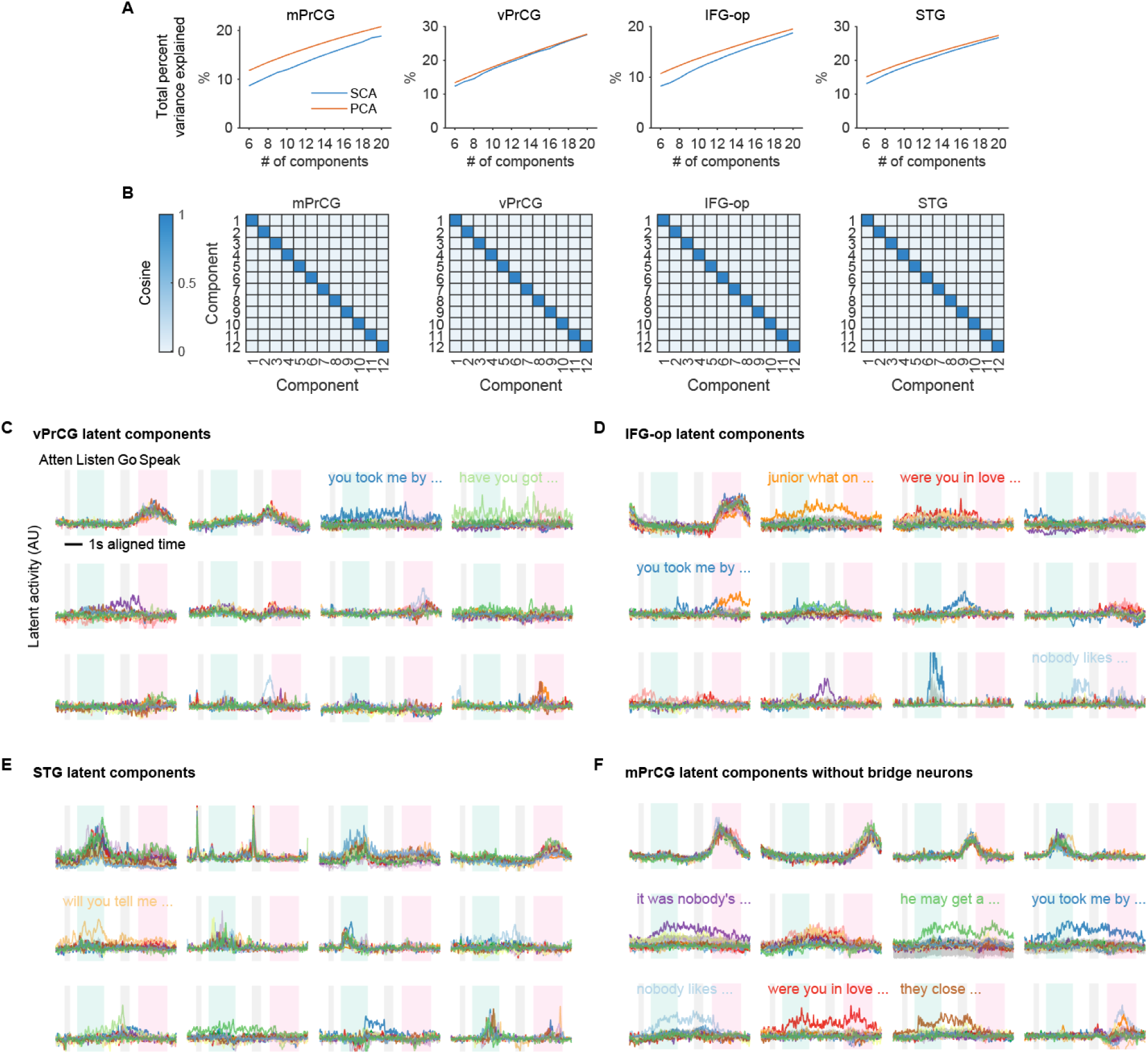
SCA latent activity across regions and conditions. (A) Comparing the total percent variance explained by SCA (blue) and PCA (red) at different numbers of components for each cortical region. (B) For each of the four regions, pairwise cosine angles show that the 12 SCA components are orthogonal. (C) Top 12 latent SCA components of the vPrCG population (n = 152 neurons). Colored traces represent the 14 different sentences. Shading around traces represents the 95% bootstrap confidence interval of projected activity using shuffled trials. In each plot, the shaded periods from left to right indicate attention cue (Atten), listening (Lisen), go cue (Go), and speaking (Speak). The text labels the sentence with significant high activation across task phases from listening to speaking. (D,E) Similar to (C) but for IFG-op (n = 329 neurons) and STG (n = 236 neurons) populations, respectively. (F) Latent SCA components of the mPrCG population excluding bridge neurons (n = 486 neurons). Same conventions as in (C).

**Figure S6.**
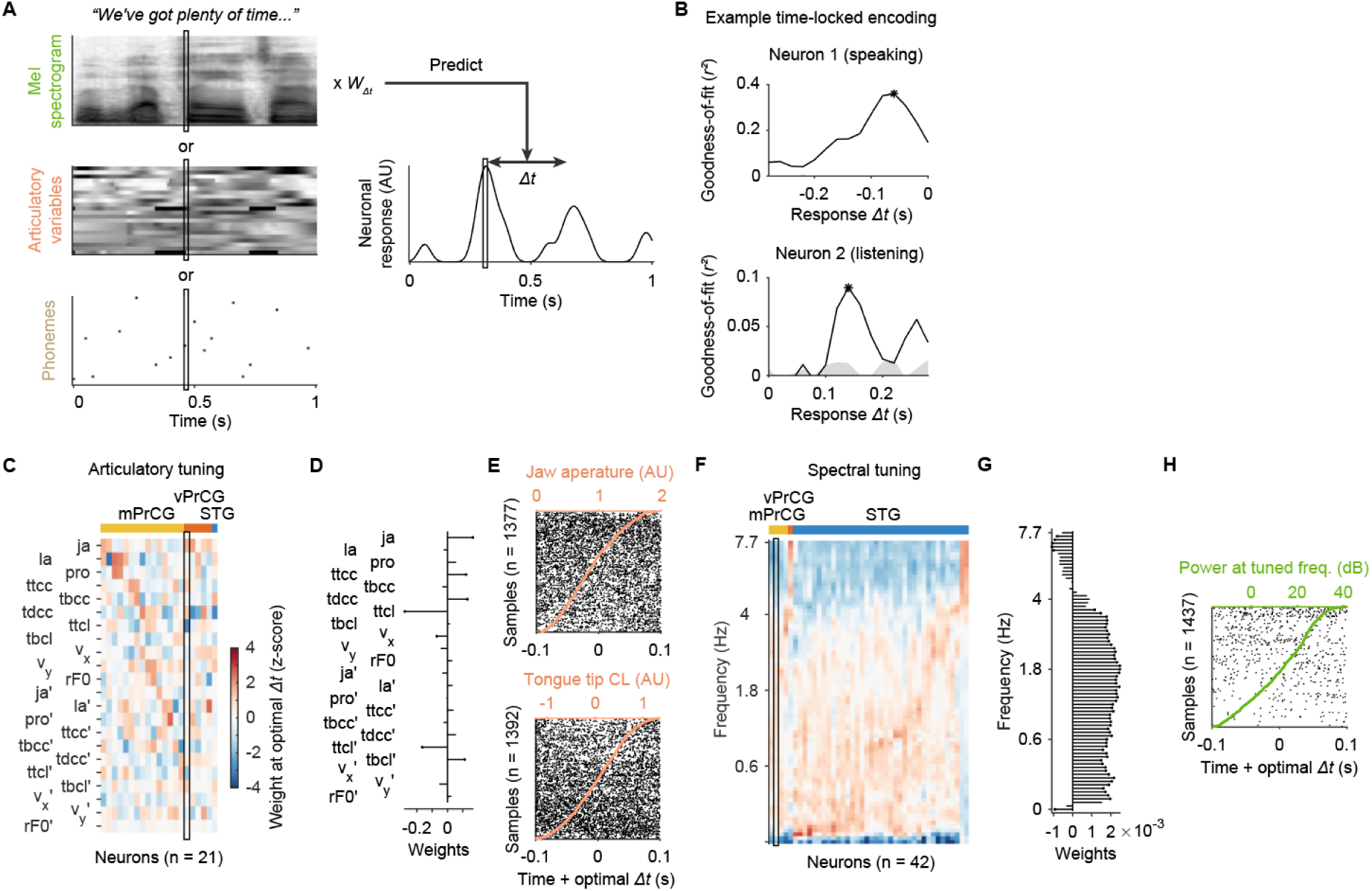
Articulatory and spectral tunings. (A) Illustration of model fitting with varied offset time (*Δt*) between speech features (left) and neuronal response (right). *W_Δt_* represents the feature weights at a given *Δt*. (B) Goodness-of-fit (*r*^2^) of example phoneme models of a speech production neuron (top) and a stimulus perception neuron (bottom) as a function of response time offset, *Δt*. Asterisks indicate the optimal encoding. Shades show the 95% confidence interval of *r*^2^ from bootstrap null models. (C) Normalized model weights across neurons (columns, groups by regions and sorted by peak weight position) that have significant articulatory encoding during speaking. The neuron highlighted in the box is expanded in (D-E). (D) Weights of the articulatory model highlighted in (C). Weights with stem heads are statistically significant. (E) Spike rasters (black) of the neuron in (D) that is modulated by tongue tip constriction location (CL) (left; orange) and jaw aperture (right; orange). (F) Similar to (C) but for neurons with significant spectral encoding during listening. (G) Weights of the spectral model highlighted in (F). Weights with stem heads are statistically significant. (H) Spike rasters (black) of the example neuron in (G) modulated by the power (green) at its most preferred spectral frequency.

**Movie 1. Neuronal responses across four cortical regions during listening, delay, initiation, and speaking phase of the delayed sentence repetition task.**

Left, two-dimensional UMAP embedding of task evoked activity of 1217 neurons (dots) across mPrCG, vPrCG, IFG-op, and STG. The dot’s color indicates the most preferred task phase of each neuron. The dot’s brightness shows the neuronal spike rate as a function of time. A wave of neuronal activity travels clockwise as a trial progresses from listening (blue; bottom left), delay (green; top), initiation (yellow; top right), and production (red; right). Upper right, sentence transcripts and task phase labels. Lower right, rasters of eight simultaneously recorded mPrCG neurons, colored by the most preferred task phase. Audio plays the microphone recording of the task performance and the simultaneous spiking sounds of u7.

**Movie 2. Mirror neuron responses to the same speech elements during listening and speaking.**

Each side of the movie shows an example mirror neuron. The top half shows responses during listening and the bottom half shows responses of the same neurons during speaking with time dynamically aligned to listening. In each half, plots from top to bottom are 1) transcribed words and a ribbon indicating the task period of listening (green) or speaking (magenta), 2) Mel spectrogram (0-8kHz) of speech sounds, 3) Spike raster (ticks) overlaid on normalized mean spike rate (trace).

**Movie 3. Sentence-selective spiking of a bridge neuron link listening and speaking across delay.**

The top and bottom each shows spiking responses during trials of a unique sentence. In each half, plots from top to bottom are 1) ribbons indicating the listening (green) and speaking (magenta) phases and the Mel spectrogram (0-8kHz) of speech sounds, 2) Spike raster (ticks), listening offsets (blue marker) and speaking onsets (red marker).

## Methods

### KEY RESOURCES TABLE

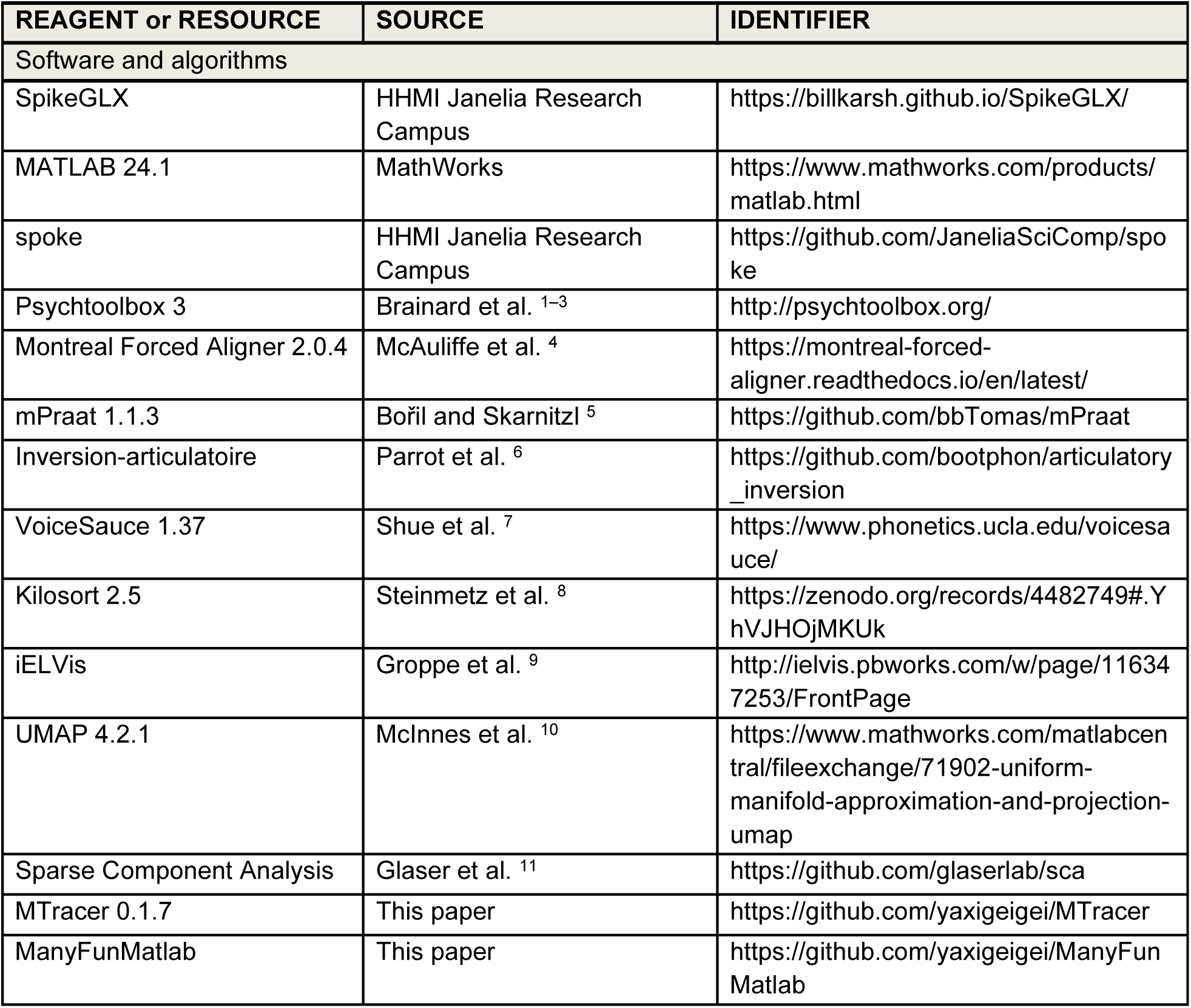

### CONTACT FOR REAGENT AND RESOURCE SHARING

Further information and requests for the data used in this study should be directed to and will be fulfilled by the Lead Contact, Dr. Edward Chang.

### EXPERIMENTAL MODEL AND SUBJECT DETAILS

#### Human subjects

The experimental protocol was approved by the UCSF Institutional Review Board. Identification and enrollment of candidate subjects proceeded in accordance with the same practices described in ^12^. Candidate subjects for intraoperative Neuropixels recording were identified among patients at UCSF scheduled to undergo awake craniotomy with speech/motor mapping for resection of a cortical lesion (see Subject Details). Informed consent was obtained in advance of the procedure by a combination of treating physicians and non-treating researchers, and participant willingness and eligibility were re-assessed at several points prior to initiation of the recording protocol.

### METHOD DETAILS

#### NP probes and acquisition system

Neuropixels recordings were performed following the methods previously described in ^12^, together with certain modifications introduced to the protocol after the cases described there which were judged to improve safety and yield of the recordings.

In advance of each recording, a Neuropixels probe, together with accompanying headstage, cabling, and custom-built tissue stabilization pedestal (Cura BioMed Clear Resin), was mounted on a micromanipulator (Narishige M-3333) which was in turn mounted to a 3-joint mounting arm (Noga NF9038CA) and clamp (Manfrotto 386BC-1). After testing functionality of all components, this entire assembly was wrapped in standard aseptic fashion and EtO-sterilized under previously validated and IRB-approved cycle parameters.

The assembly was unwrapped and positioned in the sterile field (see Intraop Procedures) and the probe connected to the acquisition system. Data were collected using standard acquisition systems ^13^ and the SpikeGLX software. The signal quality was assessed in real time for single-unit waveform quality and for speech responses using “spoke”, an online spike-raster display time-aligned to listening and speaking onsets using an custom event detection program running on an external microcontroller (Teensy 3.5).

#### Intraop procedures

Each participant was secured with a Mayfield skull clamp. Craniotomy proceeded in standard fashion and once the brain surface was exposed, a combination of neuronavigation and stimulation mapping was performed to confirm planned resection boundaries. A compatible C-clamp was then attached to the skull clamp, and in turn, a primary articulating bar was attached to the C-clamp (all items contained in Integra Brain Retractor System A2012). The clamp from the micromanipulator assembly (see NP probes and acquisition system) was then attached to the primary articulating bar. The articulating arms were positioned to place the micromanipulator above the target insertion site, the coarse X-stage of the micromanipulator was used to bring the tissue stabilizer into gentle contact with the cortical surface surrounding the target insertion site, and the fine X-stage of the micromanipulator was used to lower the Neuropixels probe through an aperture in the stabilizer to a target depth of 6 to 8 mm from the brain surface, at a rate of 50– 75 μm/sec. Insertion trajectory was approximately perpendicular to the surface at the apex of the targeted gyrus.

#### Recording site localization

The position of each Neuropixels insertion site on the cortical surface was photographed intraoperatively and recorded in the intraoperative neuronavigation system. These two records were checked for agreement and jointly used to guide manual identification of the insertion point on each subject’s preoperative MRI, a 3D reconstruction of each subject’s cortical surface (when available), and the most appropriate representative of this point on a template brain. The intracortical trajectory at the insertion site was identified using histology (when available).

#### Cortical surface localization

To locate the cortical depth of each neuron, one first needs to know where the cortical surface is located on the probe. Similar to previous studies ^12,14^, we take advantage of the distinct patterns of local field potential (LFP) in different media (tissue, saline, or air) to identify the cortical surface at the interface between the tissue and the saline added above, or the tissue and the air in the absence of saline. Specifically, a heatmap of spatiotemporal LFP signals (low passed at 50Hz) was plotted for each probe where the x-axis is recording time and the y-axis is channels ordered from tip to base. From the top going down the probe, the LFP map first shows a transition from high intensity signals to low spatial frequency low intensity signals. This indicates the air-saline interface. Further down, there is a second transition from the low spatial frequency low intensity signals to higher spatial frequency medium intensity signals. This indicates the saline-tissue interface. When saline is not present or very thin, the two transitions would merge into one. These patterns are most pronounced during the probe advancement or retraction for they cause more movement artifacts that contribute to the patterns. The spike map (a spatiotemporal scatter plot of spikes shaded by spike amplitudes) can also help to corroborate observations in the LFP map. The visualizations and measurements above were performed in MTracer, a custom MATLAB app designed to facilitate inspection and curation of Neuropixels recordings.

To validate the cortical surface localization, we examined whether spike waveforms vary as a function of the cortical depth (i.e. distance to surface) as previously reported ^15^. Indeed, we found consistent evidence that putative pyramidal cells (see the section “Classification of spike waveforms” below for details) with strong backpropagation into apical dendrites are predominantly located in the deep cortex (Figure S1H).

#### Motion correction

Intra-operative Neuropixels recordings face challenges of the relative movements between probes and neurons in tissue ^12,14^. Though the mechanical stabilization avoided major movements from the source, residual motions may still degrade the quality of the later cluster isolation. We used two post-hoc correction methods to maximally restore the stability of recorded signals. The first is the built-in motion correction algorithm of Kilosort 2.5. If this algorithm fails to correct the raw voltage timeseries, we will first apply a custom motion estimation and correction method to the raw data with manual intervention.

This custom method follows a similar rationale as previous work ^14^ where trajectories of motion are traced out based on salient spatiotemporal patterns in the signal. Here, we made several changes and improvements. First, since the reduced artifacts in LFP after mechanical stabilization no longer reflect clear movement patterns, we used trains of high SNR spikes at different cortical depths as landmarks to trace out the motion trajectories using MTracer. This procedure has been used in our previous work for quantifying the motions ^12^. Second, the cortex is not treated as a rigid body. Instead, a linear or nonlinear (modified Akima interpolation) scaling function was fitted between the magnitudes of motion trajectories (highpass filtered at 0.1Hz and/or the residual drift) and the cortical depths of these trajectories. Missing time ranges of a trajectory during a recording can be extrapolated using this motion scaling function and other trajectories that are present in those ranges. Lastly, a spatiotemporal displacement field was created from all the trajectories for the entire recording. This field was then used to reverse shift both the AP and LFP bands of the data to restore the stability of signals.

#### Preprocessing, spike sorting, and cluster curation

Raw voltage timeseries were preprocessed using the official CatGT toolbox with common average referencing and transient artifact removal. Spike sorting was then performed using Kilosort 2.5 with default parameters (e.g. 300 Hz highpass filtering). The automatically generated clusters were manually curated in MTracer with operations such as merging, cutting, setting clusters as noise, and etc. We implemented cluster curation functions in recent versions of MTracer that feature additional visualizations for curators to examine a cluster’s stability and relationships with neighboring clusters in 3-dimensions. Compared to other curation tools, MTracer is optimized to handle clusters in an environment where a unit’s spatial separability is a top factor of the quality control.

The curation was performed with an aim to isolate putative single-units. For a cluster that passed the curation, its final unit type was determined using both the thresholds of the contamination rate (20%) and the refractory period (1.5 ms) violation rate (1.5%), below which is considered a true single-unit, otherwise a multi-unit (Figure S1B).

#### Task

A trial of the standard delayed sentence repetition task starts with a beep (392 Hz, 0.3 s) drawing participants’ attention to listen to an upcoming auditory sentence stimulus. A variable delay (drawn from exponential distribution with 1s mean and capped at 5s) is introduced after the stimulus where participants maintain the perceived sentence in working memory. After the delay, a second beep (330 Hz, 0.3 s) prompts the participant to initiate and utter the same sentence. The standard task uses 14 unique sentences selected from the TIMIT corpus ^16^. A recording aims to complete a total of 72 trials where 4 of the 14 sentences each occurs 8 times and the other 10 sentences each occurs 4 times. Different sentences are interleaved and span evenly across the recording. This stimulus set strikes a balance where different sentences can sufficiently sample the phonetic space of natural speech (Figure S1I) and multiple occurrences of each unique sentence ensure robustness in estimating neuronal responses and facilitate data driven discoveries from response patterns. Early versions of the task explored the balance between the number of unique sentences and the number of repeats. Among the 20 recordings included in this study, one recording used the first version of the task which contains 29 unique sentences, two used the second version which contains 23 unique sentences, and the other 17 all used the standard version. Both of the early versions contain the 14 sentences in the final standard version.

#### Speech feature extraction

The labels and timestamps of words and phonemes were extracted using Montreal Forced Aligner (MFA v2.0.4) ^4^ with a pre-trained English model (from MFA v1). MFA requires audio waveforms and corresponding transcripts as input, and uses a dictionary to parse words into constituent phonemes. For stimuli, the audio waveforms were captured from the same speaker channel that delivered stimulation to participants. The transcripts were directly obtained from the corpse. For produced speech, the audio waveforms were recorded from the microphone. The transcripts were initialized to be the same as those for stimuli and then reviewed and updated by human curators to match the actual utterances of the participants. We used the Librispeech lexicon ^17^ as the base dictionary. If a participant utters a sound that cannot be transcribed to a real word, a new word and its phonetic composition are defined and added to the lexicon.

Similar to the procedures in previous work ^18^, the articulatory kinematic trajectories (AKTs) were estimated from audios but using an updated acoustic-articulatory inversion model ^6^. The model output contains 18 variables from which 4 additional variables are derived. Detailed description of these variables and their derivation can be found in the supplemental table 2.

The AKTs described above only capture the supralaryngeal articulation. To complement that, we use pitch as a proxy for the laryngeal control as in the previous work ^18^. A few modifications and improvements are made here. First, pitch was extracted using the VoiceSauce suite ^19^ applying the STRAIGHT algorithm ^20^. This algorithm is more robust to creaky voices from some of the participants. Second, the raw pitch timeseries during voicing periods were piecewise denoised using Savitzky-Golay smoothing (2nd order polynomials over 0.1 s frame length). Third, relative pitch was derived by normalizing the denoised absolute pitch as performed in previous study ^21^. The rate of change in relative pitch was computed by taking a discrete time derivative.

#### Dynamic time alignment

Participants spoke at different speeds. Each individual can also say the same sentences slightly differently across repeats. Moreover, some may make occasional mistakes and do not always repeat the sentences verbatim. However, many analyses and visualizations require data with temporally aligned speech and other task events. Given the multiple task phases and the sequence nature of the sentences, aligning time to each event of interest separately is unfeasible. Dynamic time alignment aims to non-linearly morph event times or timeseries such that all the events or features are transformed to a common time frame.

To align speech, we take advantage of the phonemic labels and times from MFA and use a modified Needleman-Wunsch algorithm to identify corresponding phonemic events. The modification includes a commonly used penalty of −1 for gap in addition to mismatch and indel. For repeated trials of each unique sentence, one trial is identified where the produced speech best matches the stimulus. This is the trial that has the highest alignment score (i.e. lowest edit distance) between phoneme sequences of the stimulus and the produced speech. We will later refer to this trial as the target trial. Next, separate alignments are computed between the target trial and each of the other trials. This produces rank-matched sequences of phoneme times for every trial. Missing or extra phonemes are accommodated. However, before downstream analyses, we exclude highly inconsistent trials with alignment scores (normalize to sequence length) less than 0.5. When we also need to align task phases together with speech, task event times (including attention cue onset and offset, stimulus offset, and go cue onset) are added to the phoneme times in temporal order.

A pair of rank-matched time sequences described above forms an interpolant that can morph other event times data (such as phonetic event times) and the timestamps of timeseries data (such as spectrograms or spike rates) from one time frame to another. Importantly, time-dependent derivations of speech features such as the rate of change in AKTs were computed first and dynamically aligned after. This avoids artificial distortion of signal magnitudes if aligning first and deriving after. Similarly, only spike rates computed from original data, but not spike times, are dynamically aligned. Spike trains are later reconstructed from the aligned spike rates by resampling with 0.1 ms bin size. The reconstructed spike trains were only used for visualization purposes, not for further analysis.

In this study, alignments were performed at three scopes based on different analysis requirements. The first, denoted as A^sent^, separately aligns the repeated trials for each unique sentence. The second, denoted as A^all^, further aligns task phases (using the task event times listed above) across all trials of the A^sent^ data. The third, denoted as A^l-s^, further aligns the produced speech (speaking) to the stimulus (listening) within each trial of the A^sent^ data (Figure S3A,B). The version each analysis used is described in the respective section below. Original data is used by default.

### QUANTIFICATION AND STATISTICAL ANALYSIS

#### Classification of spike waveforms

Spatiotemporal waveforms (STW) of single action potentials are extracted across 10 channels (vertically spanning 200 microns) and 2.7 ms. The extraction is centered at the detected spike time and the channel closest to the STW centroid. The centroid is computed by the mean value of channel positions weighted by the waveform powers (sum-of-squares) at those positions. For each unit, we compute the median STW and report the error (Figure S1D) by converting mean absolute deviation to standard deviation for robustness to outlier waveforms.

The identification of spike waveform types is done by nonlinear dimensionality reduction across all units followed by clustering. Specifically, UMAP ^10^ was used to reduce the 820 dimensions of median STWs to 2 dimensions (Figure S1E). This resulted in four clear aggregates of samples which were unambiguously clustered using a Gaussian mixture model with a component number of four. The average STWs of these clusters show that they are putative fast-spiking cells (FS), putative pyramidal cells (Pyr), and cells with narrow (PosN) or wide (PosW) positive spikes (Figure S1F). The validity of FS vs Pyr were further demonstrated by their distinct distributions of trough-to-peak time (Figure S1G).

#### Task responsiveness

The responsiveness of a unit at a given task phase is determined by testing whether the *N* mean spike rates during that task phase is significantly different (two-sided paired Wilcoxon signed rank test with *p* < 0.05) from the *N* mean spike rates during the baseline period, where *N* is the number of trials. For each trial, the time windows to calculate the mean spike rates are defined as follows. 1) Baseline: −0.3 to 0 s from the attention cue onset. 2) Attention: from attention cue onset to stimulus onset. 3) Listening: from stimulus onset to offset. 4) Delay: from stimulus offset plus 0.2 s to go cue onset. The added 0.2 s is a grace period to minimize confounds from unfinished responses to stimulus. Trials with this window duration less than 0.2 s are excluded from analysis. 5) Initiation: from go cue onset to production onset minus 0.2 s. The subtracted 0.2 s is a grace period to minimize confounds from early production activity. 6) Speaking: from production onset to offset.

Since a unit may respond only to a subset of trials due to speech selectivity, testing the responsiveness with all trials can dilute the selective responses and lead to false negatives. Therefore, we additionally test responsiveness using trials for each of the four most repeated sentences. A unit is considered responsive in a given task phase if it is responsive across all trials or to any of the four sentences in that phase. A unit is considered task-responsive, if it is responsive in any of the task phases including attention, listening, delay, initiation, and speaking.

#### Sentence selectivity

The sentence selectivity of a unit at a given task phase is determined using the Kruskal-Wallis test on mean spike rates during that task phase from the 4 most repeated sentences or from the standard set of 14 sentences. The time windows where mean spike rates were computed are the same as described above on task responsiveness. The strength of sentence selectivity is quantified using the Chi-squared test statistic.

#### Spike rate timeseries

Timeseries of spike rates are computed by first binning the raw spike trains at 2.5 ms and then smoothed with a 15 ms SD Gaussian kernel. Analyses in all the following sections use these spike rates or dynamically aligned versions of them unless specified otherwise.

The heatmaps of evoked activity in Figures 1C-F and S2E-H used A^all^ data such that fully aligned trials of different sentences can be averaged. For each unit, the average activity during the baseline period was subtracted to highlight the task modulation. The evoked activity is then normalized to its peak value. Single-trial responses in Figures S2I-L also used A^all^ data. Timeseries of spike rates in Figures 2, 3, 6, S3, and S4 used A^sent^ data where speech is aligned across trials with the same sentences.

#### NMF clustering

To reveal the major temporal motifs of neuronal responses in an unsupervised way, we used non-negative matrix factorization (NMF) to cluster task responsive neurons into groups, separately for each cortical region. The input to the clustering was similar to a heatmap of neuronal spike rates shown in Figure 1C-F but without the baseline subtraction. We performed the clustering with a range of cluster numbers, *k*, from 1 to 15, and computed the gap score for each *k*. The *k* value associated with the highest gap score indicates the optimal tradeoff between goodness-of-fit and model complexity. However, NMF can be sensitive to small variations in input and initialization. Therefore, we bootstrapped the above procedure 100 times where 90% of randomly sampled neurons were used in each iteration. This produced 100 *k* values associated with the highest gap scores, and we took the median as the optimal *k* for each region.

#### Movies

In Movie 1, UMAP ^10^ was used to reduce the dimensionality of the task evoked activity (as shown in Figures 1C-F, concatenated across neurons) to two dimensions. Each neuron was plotted as a dot in the two dimensional embedding space (Movie 1; left). The dot’s color indicates the most preferred task phase. The dot’s brightness shows the neuronal spike rate as a function of time. To make the activity of all 1217 neurons in sync with the audio playback, the A^sent^ data of all 20 recordings was further aligned to the target trials (see definition in the section “Dynamic time alignment”) of the example recording. To aid the visualization of task phase progression in the embedding plot, the timeseries of task evoked activity were smoothed with a 100 ms SD Gaussian kernel. To avoid color saturation, the activity was normalized to the peak response plus an extra 1 spike/sec.

#### Linker models

To assess the relationship between the perception and production activity of a neuron, we applied a linear model that predicts spike rates during speaking from spike rates during listening using the A^l-s^ data where the speech in speaking and listening are dynamically aligned (Figures S3A,B):

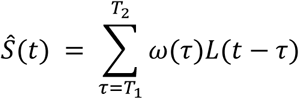

where *L*(*t* − *τ*) is the spike rate at time *t* − *τ* during listening, and *ω*(*τ*) is the linear kernel weight at time lag *τ*, and *S*^(*t*) is the estimated spike rate at time *t* during speaking. The time window for the weights ranges from *T*_1_ = −0.4 s to *T*_2_ = 0.4 s.

To reduce noise in the input, the spike rates in *L* and *S* were averaged across sentence repeats with the SEM across the repeats subtracted from the means. This error-discounted average retains the mean spike rates when a neuron fires consistently across trials but discredits the mean spike rates when large variabilities were introduced by outlier activations (e.g. bursting) in a small fraction of trials) or units drifting out halfway through the recording.

To avoid overfitting, we fitted the model with Tikhonov regularization:

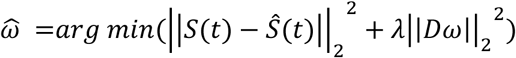

where *S*(*t*) and *S*^(*t*) were the observed and predicted spike rate for speaking at time *t*, *λ* is the regularization hyperparameter optimized for individual neurons. Specifically, we chose a difference matrix *D* as the Tikhonov matrix to enforce the temporal smoothness of the linear kernel *ω*(*τ*), which was defined as:

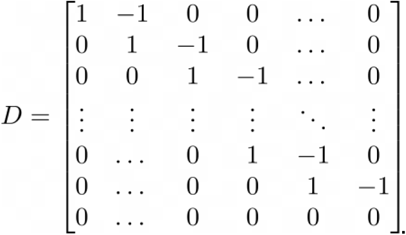

Thus, this regularization term ||*Dω*||_2_^2^ penalizes differences between the adjacent time points in the linear kernel *ω*(*τ*). The optimal *λ* was searched over a logarithmically spanned grid of 20 values from 1 to 100, and the *λ* that produces the minimal MSE across the prediction errors from 10-fold cross-validation was chosen.

The performance of this linear model was measured using the cross-validated *r*^2^. To evaluate the statistical significance, we first bootstrap sampled (n = 100 iterations) a null distribution of *r*^2^ using randomly and circularly shifted spike rates of *S* ^22^. The p-value is extracted (one-sided) from the CDF of Gumbel function fitted to the null. A model is significant if the *p*-value is less than 0.05.

The hierarchical clustering was performed across all significant models. Cosine between kernel weight vectors was used as the distance metric. Linkage was determined based on group average (MATLAB function linkage with option “average”). The order of leaves is rearranged for better visualization by maximizing the sum of similarity between adjacent leaves.

#### Linking positions

Mirror, bridge, and auditory neurons respond to specific speech content at specific sentence positions. We refer to these positions as linking positions. To algorithmically identify the linking positions for each neuron, we first quantified how well the predicted spike rates for speaking by its model matches the true spike rates.

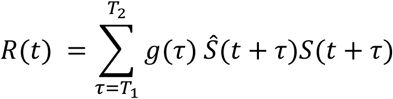

where *S*(*t*) and *S*^(*t*) were the observed and predicted spike rate for speaking at time *t*. *g*(*τ*) is a Gaussian smoothing kernel with 0.1s SD and a time range from *T*_1_ = −0.3 s to *T*_2_ = 0.3 s. We identified candidate linking positions located at peaks in *R*(*t*) that were greater than 10% of the maximum peak value. This threshold helped to remove small peaks from noise and poor matching. In addition, a valid position should have neuronal responses specific to the linked speech content. To evaluate the specificity, we tested whether the mean spike rate computed from the *N* trials of the linked sentence is significantly higher than the mean spike rate computed from *N* randomly selected trials (> 95% bootstrap confidence interval) at each candidate position. Only those with significant content-selective responses during listening and/or speaking were considered as linking positions.

#### Sentence decoding

To quantify the speech information carried by the rate code of mirror and bridge neurons and how stable the code is across time, we performed cross-time decoding of sentence identity. For each group of neurons, we first constructed a pseudopopulation with neurons pooled from different recordings in a procedure similar to previous work ^23^. We used A^all^ data where task phases and speech content across sentences were aligned. Spike rates of each neuron were de-trended, smoothed using a 0.2 s window moving average, and downsample to 10 samples/sec. For decoding, we employed a support vector machine with a linear kernel (using MATLAB fitcecoc function) to classify sentence identity from population spike rates. Only the four most repeated sentences were included as classes for they have sufficient numbers of trials. In each recording and for each sentence, we randomly set aside one trial for leave-one-out cross-validation. The remaining training trials were evenly resampled to 7 repeats (i.e. the target number of 8 repeats minus the 1 left out) such that a consistent number of trials can be concatenated across recordings. All possible train-test combinations were computed. The cross-validated accuracy was computed from 8 randomly chosen train-test combinations. This was repeated 250 times to construct a distribution of classification accuracy from which confidence intervals were estimated. To test whether a given decoding performance was significantly better than chance, we performed the same procedure as described above to compute the confidence intervals of chance accuracy with shuffled sentence labels, and compared if the observed and chance confidence intervals do not overlap.

#### Probability distribution of neurons along cortical depth

The distributions of neurons as a function of cortical depth were fitted using a non-parametric kernel smoothing method (MATLAB pdist function with a normal distribution kernel and 500 µm bandwidth). We measured how many times more likely to observe neurons in a group of interest relative to chance (Figure 4G). It is computed using the distribution of the group divided by the distribution of all task responsive neurons.

#### Sparse component analysis

The Sparse Component Analysis (SCA) ^11^ was performed separately on neuronal populations pooled in each cortical region. Using A^all^ data, an *N*-by-*T_s_* matrix of trial-averaged spikes rates were computed for each of the 14 unique sentences. *N* is the number of neurons in a given region and *T_s_* is the number of time points. The 14 matrices were concatenated into one along the time axis with a total of *T_cat_* (i.e. 14 × *T_s_*) time points. This concatenated matrix was z-scored and used as input to SCA (and PCA) to generate a lower-dimensional *Z*-by-*T_cat_* matrix where *Z* is the number of latent components.

All SCA runs used default hyperparameters from the official software package. There are two variations in model fitting. One directly imposes orthogonality constraints. The other induces orthogonality using regularization. The default method uses the latter which yielded components that are effectively orthogonal in our results (Figure S5B). It was also about an order of magnitude faster to compute. The 12 components from SCA explained 91 ± 6% (mean ± SD; n = 4 regions) of the variance that can be maximally explained by a linear dimensionality reduction method (i.e. PCA) using the same number of components (Figure S5A).

The *Z*-by-*T_cat_* matrix can be split into 14 *Z*-by-*T_s_* matrices for the different sentences. We use *v_z,s_* to denote the timeseries from the *z*-th component and the *s*-th sentence.

The criteria that determine whether a *v_z,s_* is sustained and sentence-selective takes both statistical significance and strength of activation into account. To test sentence selectivity, we first shuffled the sentence labels of the trials and computed *N*-by-*T_s_* “sentence-null” matrices of trial-averaged spikes rates. Next, these matrices were z-scored with the same factors and projected to the same latent subspace as the unshuffled data. This process was iterated 50 times and the 95% confidence interval (CI) of the sentence-null activation was computed. To examine the relative strength of activation, we tested whether each time point in *v_z,s_* is an outlier (MATLAB isoutlier function with default options) across *v_z,1_*, *v_z,2_*, …, *v_z,13_*, *v_z,14_*. A *v_z,s_* is considered sustained and sentence-selective if outlier elements go beyond the CI for at least a continuous 200 ms in each of the listening, delay, initiation, and speaking phase.

Each data point in Figure 5C quantifies the span and relative SS (sum-of-squares) of a *v_z,s_*. The span is the fraction of *v_z,s_* that is activated 3 standard deviations above the baseline (from 0.3 to 0 s before listening cue onset). The relative SS (*RSS*) of *v_z,s_* is the fraction of SS from *v_z,s_* out of total SS from the 14 sentences for a given component. Formally,

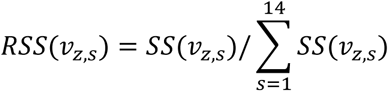

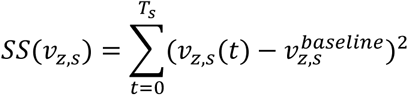

SCA produces a *Z*-by-*N* matrix of loadings, *W_n,z_*, which describes the contribution of the *n*-th neuron to the *z*-th component. To evaluate the contribution of mirror or bridge neurons, we first found the linked sentences which were given once the linking positions are identified (described in the “Linking positions” section), and focused on the latent components that show selective activation to these sentences (described above).

#### Speech feature encoding models

To understand the neuronal encoding of speech, we fitted linear encoding models using spectral, articulator kinematic, or phonemic features to predict spike rates of a unit.

Spectral features are Mel-spectrogram powers at 80 frequency bins between 0 and 8 kHz. The estimated articulator kinematic trajectories (AKTs) include variables describing lip, tongue, jaw, and velum configurations ^6^, and relative pitch as a proxy for laryngeal control ^18^. The rate of change in these variables are also included. Phonemic features include 43 unique phonemes from the stimulus set with samples defined at phoneme onsets. Detailed description of each variable is in the supplemental table 2.

For each unit, separate encoding models were fitted at a range of time offsets between features and neuronal responses. In each model:

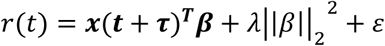

where *r*(*t*) is the spike rate at time *t*, ***x*** is a vector of feature values at time ***t*** plus a given time offset *τ*, *β* is a vector of feature weights, *λ* is the hyperparameter for L2 regularization on *β*, and *ɛ* represents the error. During listening, neurons respond to auditory input with a delay, and we use a range of *τ* from −0.4 to 0.1 s with 0.02 s steps. During speaking, we focused on neurons which signaled motor activity preceding the observed utterances, thus the *τ* goes from −0.1 to 0.4 s with 0.02 s steps. The regularization strength *λ* is zero for phonemic features since phonemes are mutually exclusive thus independent. The *λ*’s for spectral and articulatory features are empirically determined at 50 and 1, respectively.

The goodness-of-fit (*r*^2^) of a model is computed using the mean *r*^2^ from 10-fold crossvalidation. To test whether a cross-validated *r*^2^ value is statistically significant, the null distribution of *r*^2^ was sampled using bootstrap where the neuronal response was circularly shifted with random time offsets prior to model fitting. We used a significance threshold at 0.05. Negative or statistically insignificant *r*^2^ values are set to zero.

Neurons that encode segmental level features should have time-locked responses. Therefore, there must exist an optimal *τ* where the *r*^2^ is maximized. A neuron is determined as feature encoding if its *r*^2^ timeseries (e.g. Figure S6B) satisfies the following criteria: 1) The maximal *r*^2^ must be in a causal range of *τ* from −0.3 to 0 s for listening and from 0 to 0.3 s for speaking. 2) To reduce spurious fits, the maximal *r*^2^ must be greater than a small threshold of 0.01. Since we do not directly compare the magnitudes of *r*^2^ between models of different feature families, the effects of the different *λ* values on *r*^2^ will not affect the results.

#### Sequence modulation

Phonemic sequence modulation quantifies how much neuronal responses to a given phoneme can vary by different phonemes at a further position. This analysis was performed on units with time-locked phoneme encoding. For each unit, we focused on the phoneme that has the highest weight in the encoding model at optimal *τ* (see last section), and computed response modulation by different 2-phoneme and 3-phoneme sequences that start with this phoneme. During listening, the sequence elements are ordered backward in time since the neuronal response at a given time point is modulated by past phonemes. During speaking, the elements are ordered forward in time since the neuronal activity at a given time point is affected by upcoming phonemes.

We define the modulation depth (*D*) as follow:

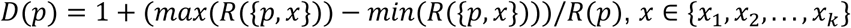

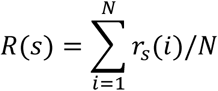

where *R*(*s*) is the neuronal response given the phonemic sequence, *s*, averaged across the *N* samples of *s*. *p* is a single phoneme or 2-phoneme sequence. *x* is a single phoneme from a set of *k* possible phonemes following *p*. {*p*, *x*} represents the concatenated sequence.

To test statistical significance, we computed the null distribution of each *D* using bootstrap where the sequence labels were randomly shuffled. All reported statistics in the main text and Figures 6J,K only include *D* values with *p* > 0.05, one-tailed (since by definition *D* ≥ 1 and the null hypothesis is *D* = 1, i.e. no modulation).

---

**Supplementary table 1.** List of participants.

| Subject | Age | Sex | Sites | Region | Hemisphere | Procedure | Pathology |
| --- | --- | --- | --- | --- | --- | --- | --- |
| NP35 | 37 | M | 1 | STG | Right | Epileptic foci resection | Subpial, subcortical, and hippocampal gliosis |
| NP38 | 38 | M | 2 | vPrCG | Left | Tumor resection | Infiltrating glioma, likely oligodendroglioma |
| NP41 | 39 | M | 1 | mPrCG | Left | Epileptic foci resection | Glial pial scar and reactive changes |
| NP43 | 67 | M | 1 | STG | Left | Tumor resection | Metastatic poorly differentiated carcinoma |
| NP44 | 29 | F | 2 | mPrCG | Left | Epileptic foci and cavernoma resection | Cavernous angioma (cavernous malformation) |
| NP45 | 70 | M | 2 | mPrCG | Right | Arteriovenous malformation and epileptic foci resection | Cavernous angioma (malformation) |
| NP46 | 67 | F | 1 | mPrCG | Left | Tumor resection | Glioblastoma, IDH-wildtype, CNS WHO grade 4 |
| NP47 | 23 | F | 1 | mPrCG | Right | Epileptic foci resection | Mild malformation of cortical development (mMCD) |
| NP50 | 69 | F | 1 | STG | Left | Epileptic foci resection | Mesial temporal sclerosis, white matter pallor |
| NP52 | 87 | M | 2 | vPrCG | Left | Tumor resection | Metastatic melanoma |
| NP53 | 29 | M | 1 | STG | Left | Epileptic foci resection | Hippocampal (mesial temporal) sclerosis |
| NP54 | 63 | F | 1 | vPrCG | Left | Tumor resection | Glioblastoma, IDH-wildtype, CNS WHO Grade 4 |
| NP56 | 27 | M | 1 | mPrCG | Right | Tumor resection | Diffuse astrocytoma, IDH-mutant, CNS WHO grade 2 |
| NP69 | 39 | M | 2 | IFG-op | Left | Tumor resection | Astrocytoma, IDH-mutant, CNS WHO Grade 2 |
| NP78 | 23 | M | 1 | IFG-op | Left | Tumor resection | Astrocytoma, IDH-mutant, CNS WHO grade 2 |

**Supplementary table 2.**
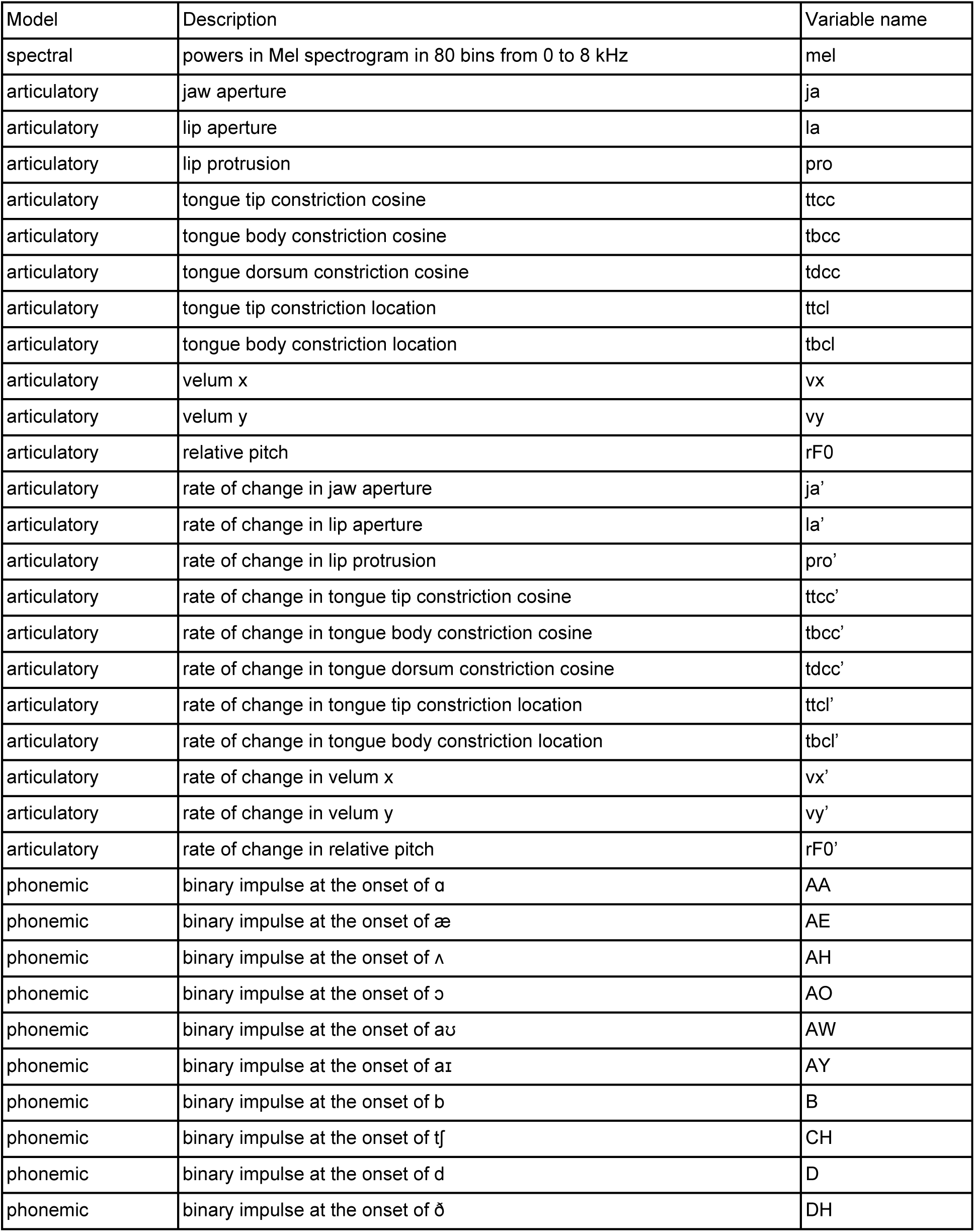
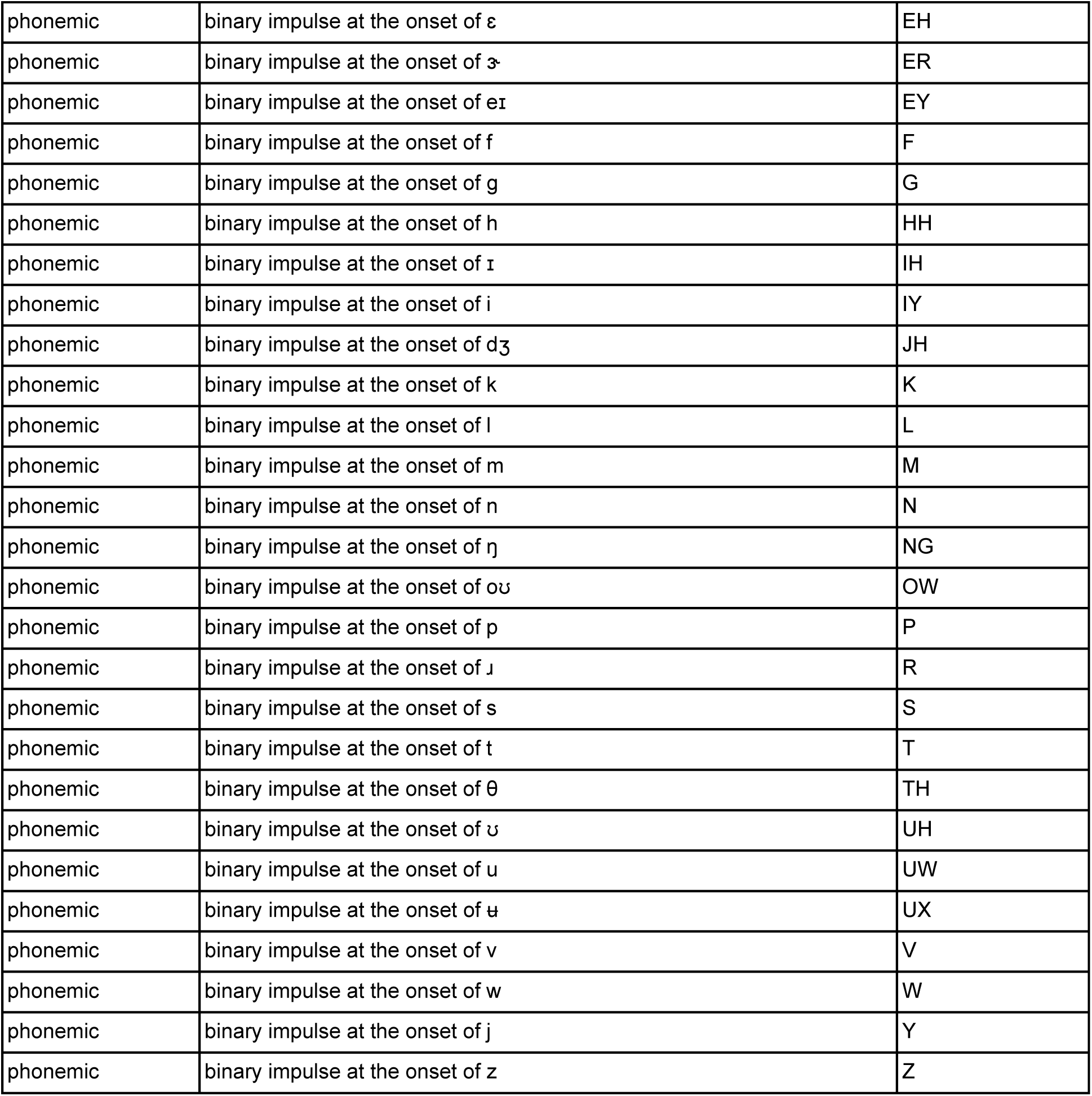
List of speech features.

## Notes

### Summary of Updates

Added Dr. Manish K. Aghi and Dr. Shawn L. Hervey-Jumper as authors who contributed to surgical operation and data collection. Updated author contributions accordingly. Fixed the ORCID of the author, Alexander B. Silva. Added acknowledgement of Dr. Adib Abla for collaboration on participant recruitment and surgical operations. Added acknowledgement to the grant from Prometheus (Fred Ehrsam).

